# Gastruloids reveal alternative morphogenetic routes for body axis elongation with distinct cytoskeletal dependencies

**DOI:** 10.1101/2025.08.04.668470

**Authors:** Guillermo Serrano Nájera, Apolline Delahaye, Benjamin J. Steventon

**Author notes:** Equal contribution (Guillermo Serrano Nájera, Apolline Delahaye).

## Abstract

Morphogenesis requires cells to integrate genetic and environmental information to pattern and shape tissues. Recent studies in vivo and in vitro have shown how alterations in environmental cues can induce alternative morphogenetic programs. However, what determines which morphogenetic strategy cells deploy in different environments remains unknown. Using gastruloids, a model for mammalian posterior body-axis elongation where environmental cues can be fully controlled in vitro, we show that substrate availability determines which force-generation strategy cells employ to build the body axis. In free-floating conditions, gastruloids self-assemble a single posterior body axis through cell-cell interactions. When plated on laminin, gastruloids acquire a flat morphology, break symmetry multiple times, and produce several independently elongating body axes through collective cell migration. On laminin, formin activity and focal adhesion-mediated traction are required for tissue elongation, yet both are functionally dispensable in free-floating conditions. Transcriptomic analysis shows that formin inhibition blocks elongation on laminin without any detectable transcriptional changes, demonstrating that these cytoskeletal components play a purely mechanical role that is only required when cells engage a substrate. Furthermore, laminin biases cells toward posterior migratory fates while preserving the Hox expression pattern. Together, these results show that mammalian body-axis elongation can proceed through different morphogenetic modes that engage distinct mechanical effectors depending on substrate availability. These findings have implications for understanding how evolution explores morphological diversity and how tissue engineering might achieve desired forms through precise environmental control.

## Introduction

Animal morphogenesis requires cells to coordinate their behaviours to build tissues and organs by integrating genetic programs with environmental signals. The interplay between these factors can result in morphogenetic plasticity: variable developmental outcomes despite identical genetic backgrounds. Recent work has demonstrated this both in vivo and in vitro, showing that geometry, mechanical properties, and chemical composition of the environment can alter morphogenetic outcomes (Chuai et al., 2023; Muncie et al., 2020; Karzbrun et al., 2021). A clear example is gastrulation, where alterations in geometry, maternal determinants, or biochemical environment can drastically change the mode of mesoderm internalisation in cnidarians, insects, and vertebrates (Serrano Nájera and Weijer, 2023). This raises a fundamental mechanistic question: when developing tissues adopt different morphogenetic strategies, to what extent does this depend on the activation of new gene-regulatory programs versus the context-dependent use of existing cellular machinery? Understanding these interactions is critical for revealing the developmental rules underlying robustness, evolvability, and synthetic morphogenesis. However, studying them in developing embryos remains challenging due to maternal constraints, system complexity, and limited experimental accessibility.Moreover, while alternative morphogenetic outcomes have been reported in vitro and in vitro, morphogenetic plasticity itself has not been the subject of direct experimental dissection.

For these reasons, morphogenetic plasticity may be better studied in vitro, where environmental conditions can be precisely controlled. Gastruloids, stem cell aggregates that recapitulate mammalian posterior body-axis elongation (Van den Brink et al., 2014; Van Den Brink et al., 2020), are ideal for studying how environmental context affects morphogenesis: they manifest a reproducible morphogenetic program, can be produced in high-throughput, and grown within a fully defined environment (Anlas and Trivedi, 2021; Morales et al., 2021; Serrano Nájera and Weijer, 2023), while reproducing key aspects of mammalian gastrulation, including derivatives of the three germ layers, epithelial-to-mesenchymal transition (EMT) hallmarks, and symmetry breaking(Van den Brink et al., 2014; Van Den Brink et al., 2020).

However, during normal mammalian gastrulation, complex cell-substrate interactions drive key morphogenetic processes including EMT and cell migration through the primitive streak (Leptin, 2005). This unfolds within a dynamically remodelling extracellular environment that begins with a laminin-rich basement membrane and progresses through sequential deposition of collagens and fibronectin (Leivo et al., 1980). Instead, standard protocols culture gastruloids as free-floating aggregates (Van den Brink et al., 2014), preventing substrate attachment and limiting morphogenetic forces to cell-cell interactions (Hashmi et al., 2022; Oriola et al., 2024; Fiuza et al., 2024; Gsell et al., 2025; McNamara et al., 2024). Alternative approaches use micropatterning to grow embryonic stem cells on matrix-coated spots of defined size and shape, producing robust concentric germ layer patterns (Warmflash et al., 2014; Morgani et al., 2018). While micropatterning enables precise control over geometry and mesoderm induction sites (Muncie et al., 2020), the confined nature and adhesive coatings restrict morphogenetic movements, preventing formation of a patterned body axis. As a result, how ECM substrates affect morphogenesis in gastruloids remains untested.

Here we show that gastruloids plated on laminin without micropatterned constraints generate patterned posterior body axes using an alternative morphogenetic mode through collective cell migration. Using quantitative morphological analysis, pharmacological perturbation, and transcriptomic profiling, we demonstrate that laminin substrates induce transcriptional changes that bias cells toward posterior migratory fates while preserving the profile of Hox gene expression. On laminin, body-axis elongation requires formin activity and focal adhesion-mediated traction, yet both are functionally dispensable in free-floating conditions. Together, these results show that body-axis elongation can proceed through different morphogenetic modes that engage distinct cytoskeletal effectors depending on substrate context.

## Results

### Mesoderm specification regulates gastruloid substrate-adhesion profile

Morphogenetic processes such as cell migration, epithelial folding, and tissue elongation require mechanical forces generated through cell-cell or cell-substrate interactions. In standard gastruloid culture, cells aggregate in ultra-repellent plastic wells that prevent substrate attachment (Van den Brink et al., 2014), creating a system where morphogenesis relies mostly on cell-cell interactions to generate forces.

Here we probed the developmental potential of gastruloids in the presence of various adherent substrates. We developed an automated imaging pipeline to quantify the dynamics of reporter gene expression, morphology, and cell migration on gastruloids cultured on ECM-coated surfaces (Fig. 1A). By day 5, gastruloids cultured on these surfaces exhibit diverse morphogenetic outcomes depending on the substrate, which differ significantly from the simple elongated shape of regular free-floating gastruloids (Fig. 1B). For instance, gastruloids growing on laminin develop multiple flat projections at their periphery and a halo of individually migrating cells, while gastruloids cultured on either fibronectin or collagens IV/I are characterized by a single elongating pole and variable degrees of cell migration surrounding them.

**Fig. 1.**
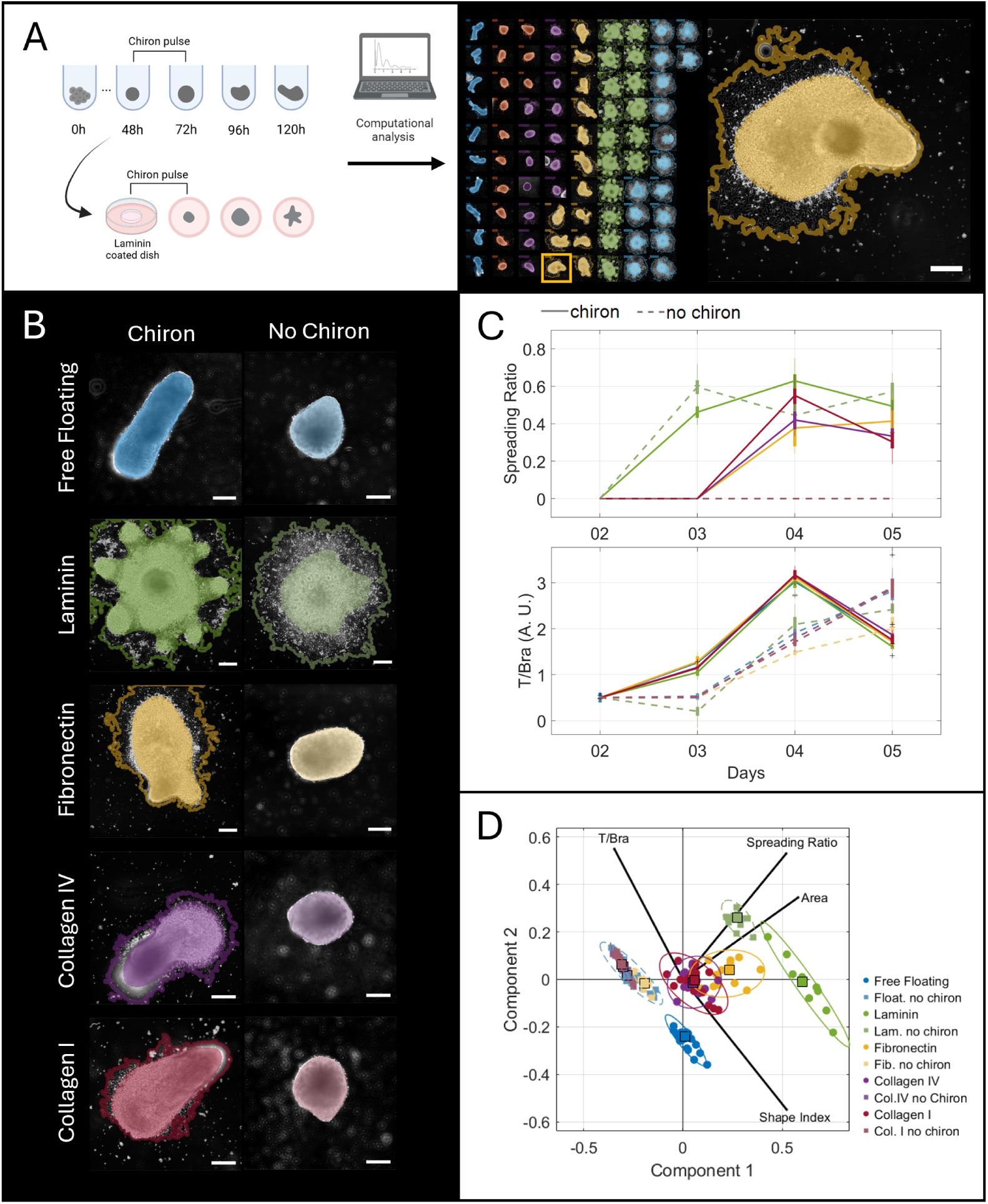
Mesoderm specification regulates gastruloid substrate-adhesion profile. A) High-throughput production (left) and analysis (centre) of gastruloids on different substrates. In the close up (right) the continuous mask covers the main body of the gastruloid while the continuous line marks the limit of the migration front. B) Gastruloids exhibit diverse morphogenetic behaviours depending on the substrate and the presence of Chiron. C) Analysis of the spreading capability (spreading ratio) of gastruloids on different substrates (top) and T/Bra expression (bottom) from day 2 to 5. D) Principal Component Analysis integrating different properties on day 5 (see Fig. S1). Colours indicate the same conditions across graphs.

Interestingly, gastruloid adhesion preferences change over time (Fig. 1D): When exposed to ECM-coated glass surfaces after aggregation (day 2), gastruloids attach exclusively to laminin, not adhering to any other coating (fibronectin, collagen I, collagen IV); however, at later stages (day 3 - day 4), gastruloids gain the ability to adhere to all tested components, suggesting a shift in the substrate adhesion capabilities following CHIR99021 (Chiron) exposure on day 2. Since Chiron stabilises mesoderm induction and mesoderm cells exhibit a distinct adhesion profile (Burdsal et al., 1993), we hypothesised that Chiron exposure could be necessary for gastruloid cells to attach to ECM proteins other than laminin. Indeed, in the absence of Chiron exposure on day 2, gastruloids failed to attach to fibronectin and collagen IV/I coated surfaces from day 3 onward, while maintaining their ability to adhere to laminin (Fig. 1B-D, Fig. S1). Notably, free-floating gastruloids not exposed to Chiron express T/Bra in a delayed fashion (Fig. 1D), but do not elongate (Fig. 1B-D, Fig. S1).

Next, we tested the effects of Matrigel, a basement membrane extract mainly composed of laminin and collagen IV (Sodek et al., 2008; Kibbey, 1994). However, despite the presence of laminin, we found that gastruloids on Matrigel did not replicate the phenotype observed on laminin. Instead, Matrigel induced a morphology similar to gastruloids exposed to fibronectin and collagens (Fig. S2A), with the same Chiron-dependent adhesion temporal profile (Fig. S2B), which suggests that other components of the extract might have a dominant effect over the laminin at this early stage.

Overall, these results demonstrate that gastruloid adhesion profiles change with their differentiation state, possibly mirroring the sequential deposition of ECM components during embryonic development. The strong affinity for laminin, independent of Chiron treatment, is consistent with laminin being the first ECM protein deposited in mammalian embryos (Cooper and MacQueen, 1983), suggesting that the cellular machinery for laminin adhesion is already present before mesoderm specification.

### Gastruloids on laminin produce multiple posterior body axes

In free-floating gastruloids, on day 4, the emergence of the axis is preceded by the formation of a T/Bra-positive pole from an initial salt-and-pepper pattern of T/Bra-positive cells (Van den Brink et al., 2014; Hashmi et al., 2022); followed by axial elongation on day 5, driven by cell-cell interactions (Gsell et al., 2025; McNamara et al., 2024; Mayran et al., 2023). We explored whether similar T/Bra-positive cell dynamics are associated with the emergence of cell streams when gastruloids are cultured on laminin. Similar to micropatterned systems (Warmflash et al., 2014; Morgani et al., 2018), we found that T/Bra-positive cells organize as a ring on the edge of flattened gastruloids on day 3, and by day 4 they coalesce into poles (Fig. 2A). Notably, T/Bra cells on laminin tend to form multiple poles rather than coalescing into a single axis as in free-floating conditions, likely due to the geometric and motility constraints imposed by adhesion to the substrate.

**Fig. 2.**
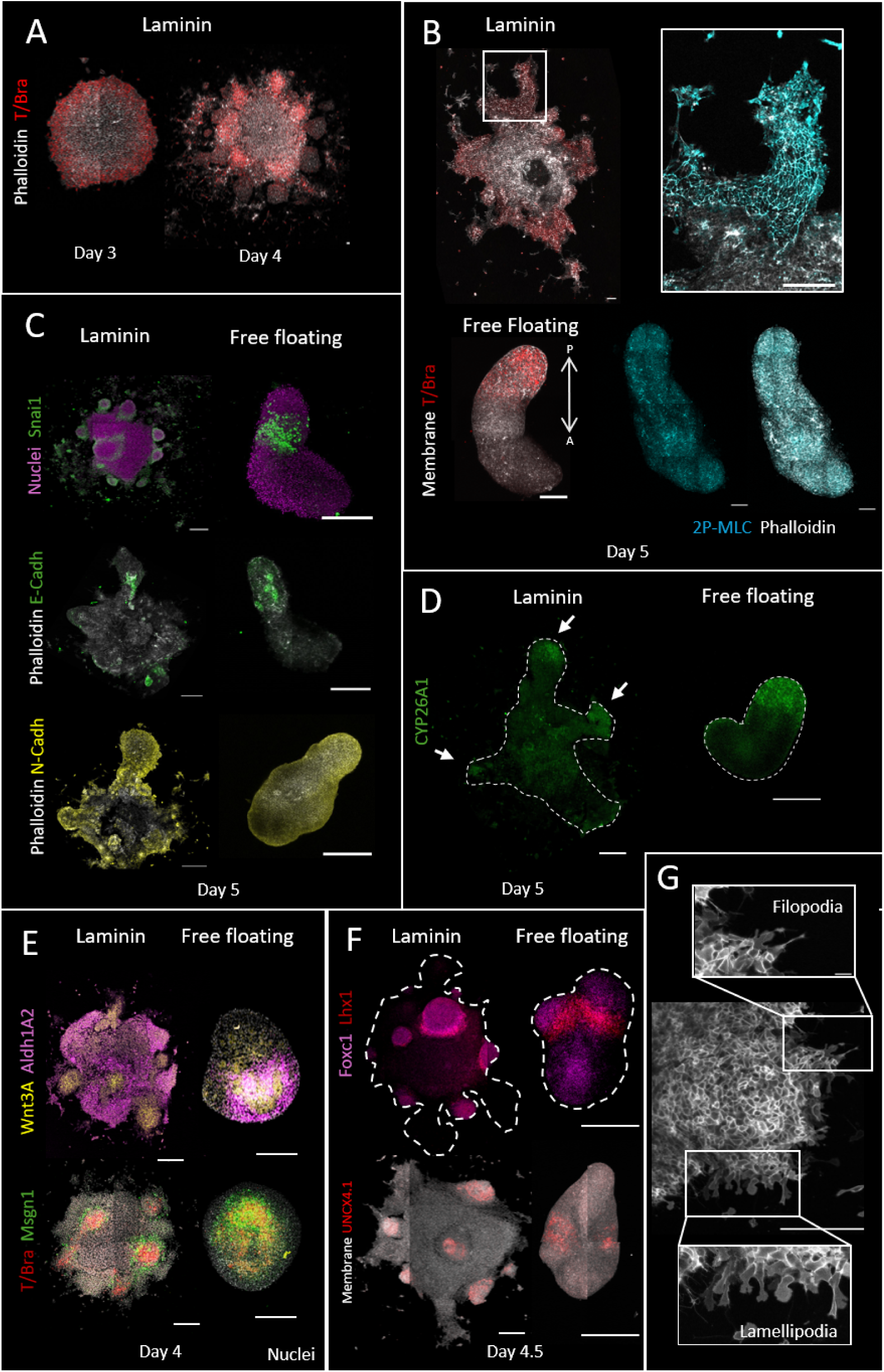
Free floating and laminin gastruloids share similar differentiation abilities. A) T/Bra immunostaining of gastruloids on laminin on days 3 and 4. B) T/Bra and double phosphorylated myosin immunostaining in free-floating and laminin gastruloids on day 5. C) EMT-associated immunostainings on day 5 in free-floating and laminin gastruloids. D) Expression of the posterior tip marker CYP26A1 in free-floating and laminin gastruloids on day 5. E) HCR for Wnt3A, Aldh1A2, T/Bra and Msgn1 in in free-floating and laminin gastruloids on day 4. F) HCR for Foxc1, Lhx1 and Uncx4.1 in free-floating and laminin gastruloids on day 4.5. G) Confocal image of a cell stream emerging in a membrane GFP gastruloid on laminin. Scale bars are 200 *µm* for the free-floating gastruloids and 500 *µm* for the laminin gastruloids.

Next, we examined the activation of proteins associated with morphogenetic movements. We observed that the emerging cell streams on laminin are associated with a clear pattern of myosin activation (Fig. 2B) where double-phosphorylated (2-P) myosin is highly enriched at cellular edges within in the elongating tip and completely absent in the main body of the gastruloid. Instead, in free-floating gastruloids, 2-P myosin shows a diffuse pattern without distinct domains (Fig. 2B). This might suggest in free-floating gastruloids multiple tissues contribute to elongation, whereas on laminin it is primarily driven by T/Bra-positive cells. However, despite these differences in its activation patterns, myosin contraction is necessary for morphogenesis in both laminin-cultured and free-floating gastruloids, as demonstrated by Rho kinase inhibition, which hinders elongation in both scenarios (Fig. S3).

Beyond myosin activation, we looked at EMT markers. EMT is characterized by the downregulation of E-cadherin and the upregulation of N-cadherin and SNAI1 (Cano et al., 2000; Lamouille et al., 2014). Since this cadherin switch regulates the elongation of conventional free-floating gastruloids (Mayran et al., 2023), we investigated whether a similar transition occurs in gastruloids undergoing axial elongation on laminin. By day 5, SNAI1 is expressed in a band at the base of the cell streams, mirroring the pattern observed in free-floating gastruloids (Fig. 2C). Similar to free-floating gastruloids, E-cadherin expression is largely lost throughout laminin-cultured gastruloids (Hashmi et al., 2022; Mayran et al., 2023; Fiuza et al., 2024), persisting only in patches, while N-cadherin is highly expressed at the outer edges of the gastruloid and within the streams (Fig. 2C). These results indicate that free-floating gastruloids and gastruloids on laminin undergo a comparable EMT.

We then tested whether gastruloids on laminin can generate the hallmarks of a patterned posterior body axis beyond T/Bra expression as in free-floating gastruloids. In free-floating gastruloids and embryos, the most posterior end expresses Cyp26a1, an enzyme involved in RA catabolism. Similarly, the distal ends of streams on laminin express Cyp26a1 (Fig. 2D, Fig. S4), indicating their patterned posterior identity. Consistent with this, we observed the expected anterior-posterior segregation pattern: Wnt3a (posterior marker) is localized within the streams, while Aldh1a2 (anterior RA synthesis marker) is expressed in the main body of the gastruloid (Fig. 2E). These findings demonstrate that cell streams possess the molecular hallmarks of posterior tissues. Lastly, we confirmed that posterior identities are necessary for cell stream formation by shifting gastruloid tissues towards more anterior fates using activin A, instead of Chiron (Dias et al., 2025). In these conditions, free-floating and laminin gastruloids fail to elongate, indicating that cell stream formation indeed depends on the presence of posterior cell fates (Fig. S5).

Furthermore, we observed enhanced spatial separation of mesodermal derivatives when gastruloids are cultured on laminin. This includes distinct localisation of early (Msgn1, Fig. 2E) and late (Foxc1, Uncx4.1, Fig. 2F) paraxial mesoderm markers, and intermediate mesoderm markers (Lhx1, Fig. 2F). Notably, gastruloids on laminin can form bulging structures that often flank the cell streams and express somite-associated markers (Foxc1, Uncx4.1, Fig. 2F) in organized clusters, contrasting with the more sparsely distributed expression observed in free-floating gastruloids (Fig. 2F). This enhanced organization supports the idea that laminin promotes somitogenesis in gastruloids, consistent with previous studies using Matrigel to induce somite-like structures (Van Den Brink et al., 2020; Veenvliet et al., 2020).

Finally, we performed live imaging using a line expressing EGFP at the cell membrane (GPI-GFP, see methods) to capture the cellular behaviours associated with cell stream elongation (Movie 1). We observed that the emergence of cell streams is a highly dynamic process, with cells migrating on the substrate while extending active filopodia and lamellipodia (Fig. 2G), actin-based subcellular structures with important roles in cell movement. Altogether, these results suggest that cell streams constitute a posterior body axis that employs different cell behaviours and myosin activation patterns than their free-floating counterparts.

### Inhibition of lamellipodia formation enhances collective cell migration on laminin

Lamellipodia are dense, two-dimensional branched network of actin filaments important for cell locomotion. To assess the role of lamellipodia in the elongation of cell streams on laminin, we used CK666 (Fig. 3), an inhibitor of the Arp2/3 complex (Hetrick et al., 2013; Nolen et al., 2009). This complex is an actin nucleator that organizes actin fibres in a branching configuration necessary for lamellipodia formation (Mullins et al., 1998; Goley and Welch, 2006). As expected, adding CK666 between days 4 and 5 reduced the formation of lamellipodia (Fig. S6) and the individual cell migration on both laminin and fibronectin (Fig. 3A-B, Fig. S7A), suggesting that lamellipodia are required for the effective directed displacement of individual cells.

**Fig. 3.**
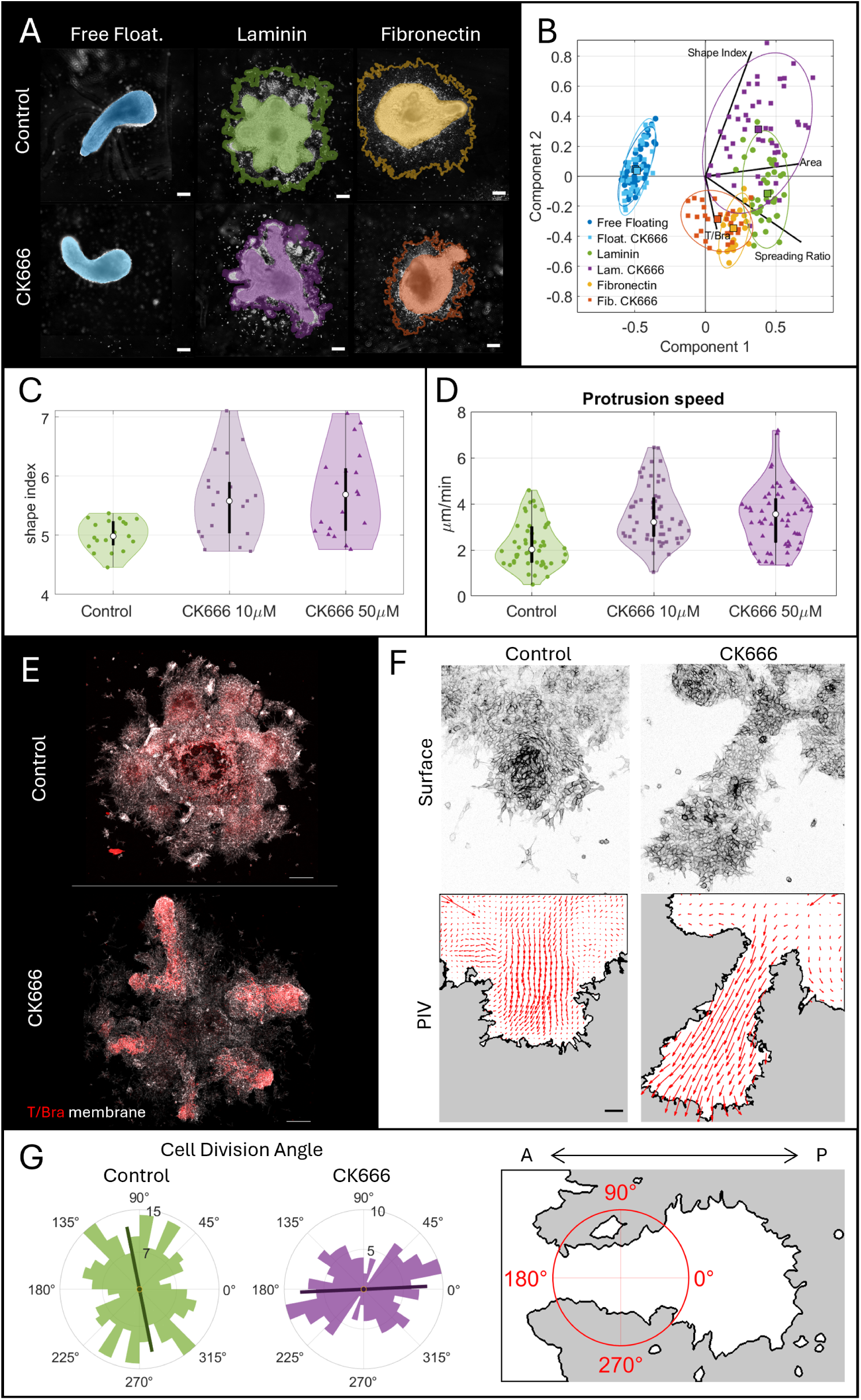
Inhibition of lamellipodia enhances elongation on laminin-seeded but not on free-floating gastruloids. A) Gastruloid morphologies on different substrates with or without CK666 50 *µM* at day 5. B) Principal Component Analysis integrating different properties on day 5 (see Fig. S7A). C) Quantification of the shape-index with different CK666 concentrations. D) Quantification of the migration speed of T/Bra positive tips E) High-resolution image demonstrating the higher elongation capabilities of laminin-seeded gastruloids treated with CK666. F) Particle image velocimetry on the developing cell streams (see Fig. S8). G) Quantification of the cell division angle with respect to axis of elongation. See supplementary material for full statistical analysis. Colours indicate the same conditions across graphs. Scale bars are 200 *µm* in A-E and 50 *µm* in F.

However, surprisingly, CK666 treatment resulted in longer cell streams protruding from gastruloids on laminin (Fig. 3C, E), with no measurable effect on the elongation of free-floating gastruloids or those seeded on fibronectin (Fig. S7). To quantify these differences, we decomposed gastruloid morphology using LOCO-EFA modes (Fig. S7), a series of unbiased bidimensional shape descriptors based on Fourier transforms (Sánchez-Corrales et al., 2018). While the first two modes describe ellipsoidal shapes, higher frequencies characterize more complex morphologies. This analysis confirmed that CK666 shifted laminin-plated gastruloids toward more complex, elongated shapes without affecting free-floating gastruloids or those on fibronectin (Fig. S7C).

Next, we sought to identify the underlying cause of the more elongated morphology. Since there was no change in the area of laminin-plated gastruloids after CK666 treatment (Fig. S7), we hypothesized that the elongation resulted from enhanced migration capabilities of cell streams (Movie 2). To test this, we quantified the migration speed of cell stream tips from gastruloids on laminin exposed to CK666 (Movie 3). Our measurements revealed that migration velocity increased, confirming that inhibition of the Arp2/3 complex enhanced their migratory capacity (Fig. 3D).

To further investigate the tissue movements, we recorded high-resolution time-lapse movies of protrusion formation using the GPI-GFP line with or without CK666. Particle image velocimetry (PIV) analysis of these movies suggested that cells move in a coherent directional manner with an increased velocity in the presence of CK666 (Fig. 3F, Fig. S8). In both conditions, we did not observe significant oriented cell intercalations once cells abandoned the main gastruloid body and entered the stream, which suggests that the formation of these streams represents a form of collective cell migration rather than convergent-extension fuelled by oriented cell intercalations. The pattern of tissue flows was not affected by the addition of CK666, except for the enhanced migration speed.

Finally, we observed that cell divisions in the stalk connecting the most distal (posterior) tip with the main gastruloid body preferentially orient perpendicular to the elongation axis (Fig. 3G). We hypothesised that lamellipodia at the perimeter of the stream (Fig. 2G) generate lateral pulling forces that bias cell division angles (Fig. 3G). These perpendicular forces are presumably counter-productive for the cell stream formation since they contribute to lateral rather than axial elongation. Strikingly, CK666 treatment caused cell division orientation to shift and statistically align with the elongation axis (Fig. 3G), suggesting that removing lamellipodia reduces lateral pulling, leaving only productive forces that promote axial elongation, increasing migration speed. On the whole, these experiments show that Arp2/3 has a context-dependent mechanical function: on laminin, lamellipodia may generate lateral forces that slow down axial elongation, whereas in free-floating conditions Arp2/3 inhibition has no effect.

### Body axis elongation in laminin has distinct cytoskeletal requirements

Having established that inhibiting lamellipodia formation enhances collective cell migration and stream elongation specifically on laminin, we next investigated the role of filopodia in this process. Filopodia are finger-like cellular protrusions composed of parallel bundles of actin filaments (Mattila and Lappalainen, 2008), which are primarily nucleated by formins (Courtemanche, 2018). To determine whether these actin-rich protrusions are required for the laminin-specific cell stream elongation (Fig. 4), we treated gastruloids with SMIFH2, a small molecule inhibitor of formin homology 2 domains (FH2) (Rizvi et al., 2009), producing a reduction in filopodial length on both laminin-plated (Fig. S6) and free-floating (Fig. S9) gastruloids.

**Fig. 4.**
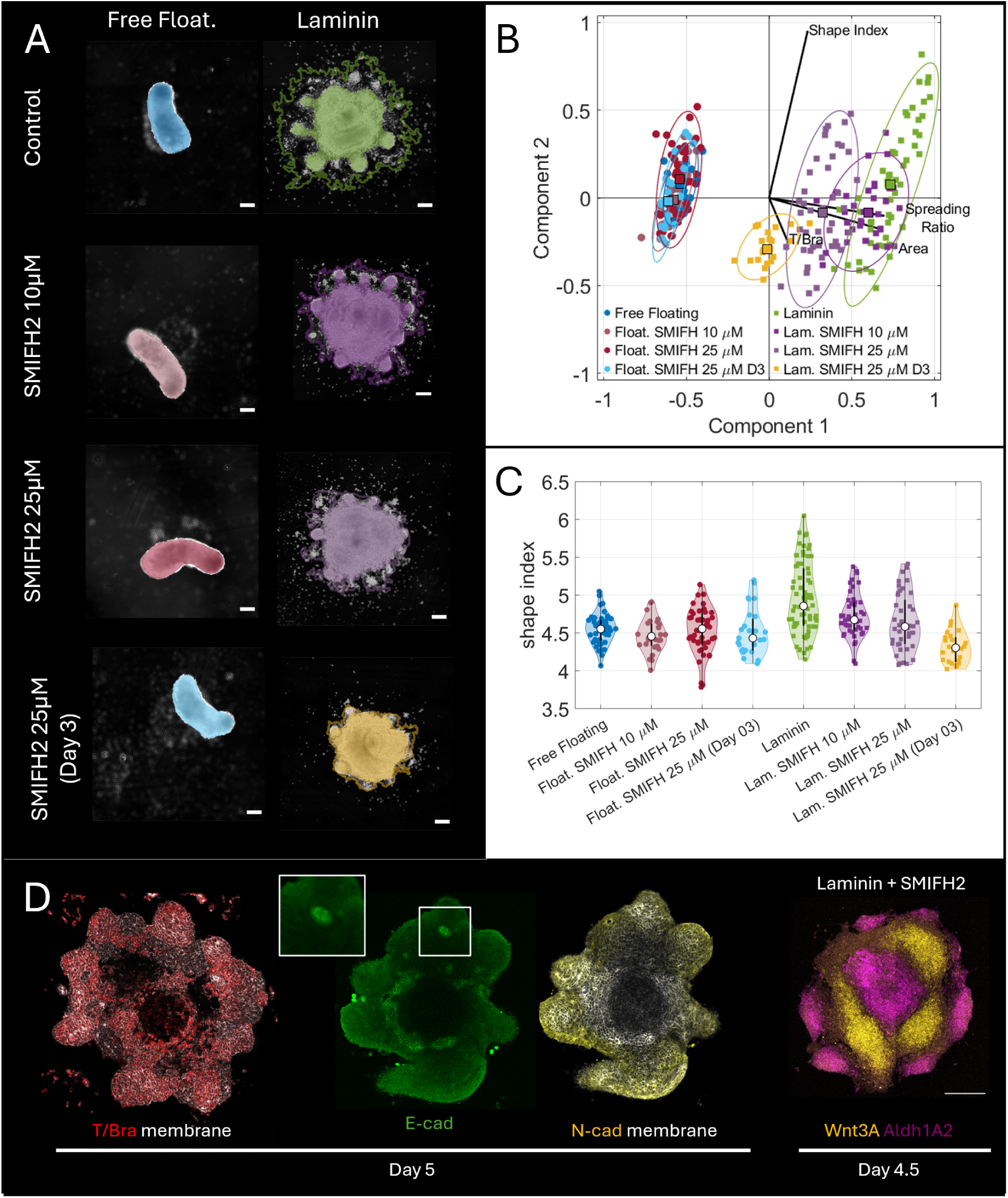
Inhibition of formin activity blocks elongation on laminin-seeded but not on free-floating gastruloids. A) Gastruloid morphologies on different substrates with or without SMIFH2 at day 5. B) Principal Component Analysis integrating different properties on day 5 (see Fig. S10A). C) Quantification of the shape-index with different SMIFH2 concentrations. D) Expression of T/Bra (reporter line), E/N-Cadherins (immunofluorescence), and Wnt3/Aldh1A2 (HCR) on laminin gastruloids treated with SMIFH2 25 *µM* at day 3 (84h). Inset shows a patch of E-Cadherin expression on the cell membrane. See supplementary material for full statistical analysis. Colours indicate the same conditions across graphs. Scale bars are 200 *µm*.

Strikingly, SMIFH2 blocked the formation of cell streams with its severity increasing with dose or exposure time, on gastruloids cultured on laminin, while having no effect on their free-floating counterparts (Fig. 4). The inhibitor prevented area expansion, individual cell migration, and cell stream formation in gastruloids cultured on laminin (Fig. S10A), but had no measurable effect on free-floating gastruloids as shown in the integrated PCA (Fig. 4B). Indeed, LOCO-EFA shape analysis demonstrated that SMIFH2 decreased the morphological complexity of laminin-plated gastruloids in a dose-dependent manner (Fig. S10B). While control gastruloids on laminin developed complex shapes characterized by higher-order LOCO-EFA modes, SMIFH2-treated gastruloids remained predominantly circular (Fig. S10C).

We did not observe significant changes in mesodermal differentiation (T/Bra), the cadherin switch (E-cad/N-cad), or the specification of anteroposterior markers (Aldh1A2/Wnt3a) upon addition of SMIFH2 at day 3.5 (84 hours), even at the highest tested concentration (25 *µM*, Fig. 4D, Fig. S10D). This suggests that filopodia inhibition does not substantially affect signalling pathways, but instead plays a critical mechanical role in generating forces on a substrate. Moreover, if filopodia were significantly involved in signalling, we would expect to see similar effects in free-floating gastruloids, which also exhibit filopodial protrusions (Fig. S9). Altogether, these results demonstrate that formin activity is essential for the formation and elongation of cell streams in laminin-seeded gastruloids (Movie 4).

### Formin activity plays a purely mechanical role specific to laminin

To distinguish whether formin inhibition blocks elongation by disrupting cell fate or by removing mechanical force, we performed a transcriptomics analysis (bulk RNA-seq) comparing the effects of laminin substrate and SMIFH2 treatment (see methods). We identified differentially expressed genes when gastruloids were plated on laminin, regardless of SMIFH2 treatment (Fig. 5A, Fig. S11). However, comparing DMSO- and SMIFH2-treated gastruloids revealed no differentially expressed genes, whether cultured under free-floating conditions or on laminin (Fig. 5A, Fig. S11, Sup. Table 1). Given the transcriptional equivalence between SMIFH2-and DMSO-treated samples we pooled them together to increase the statistical power of the analysis (n = 6 replicates per condition; see methods), and performed differential expression analysis on the combined data (Fig. 5B, Sup. Table 1). These results indicate that formin inhibition blocks axis elongation mechanically, without affecting cell fate or gene expression.

**Fig. 5.**
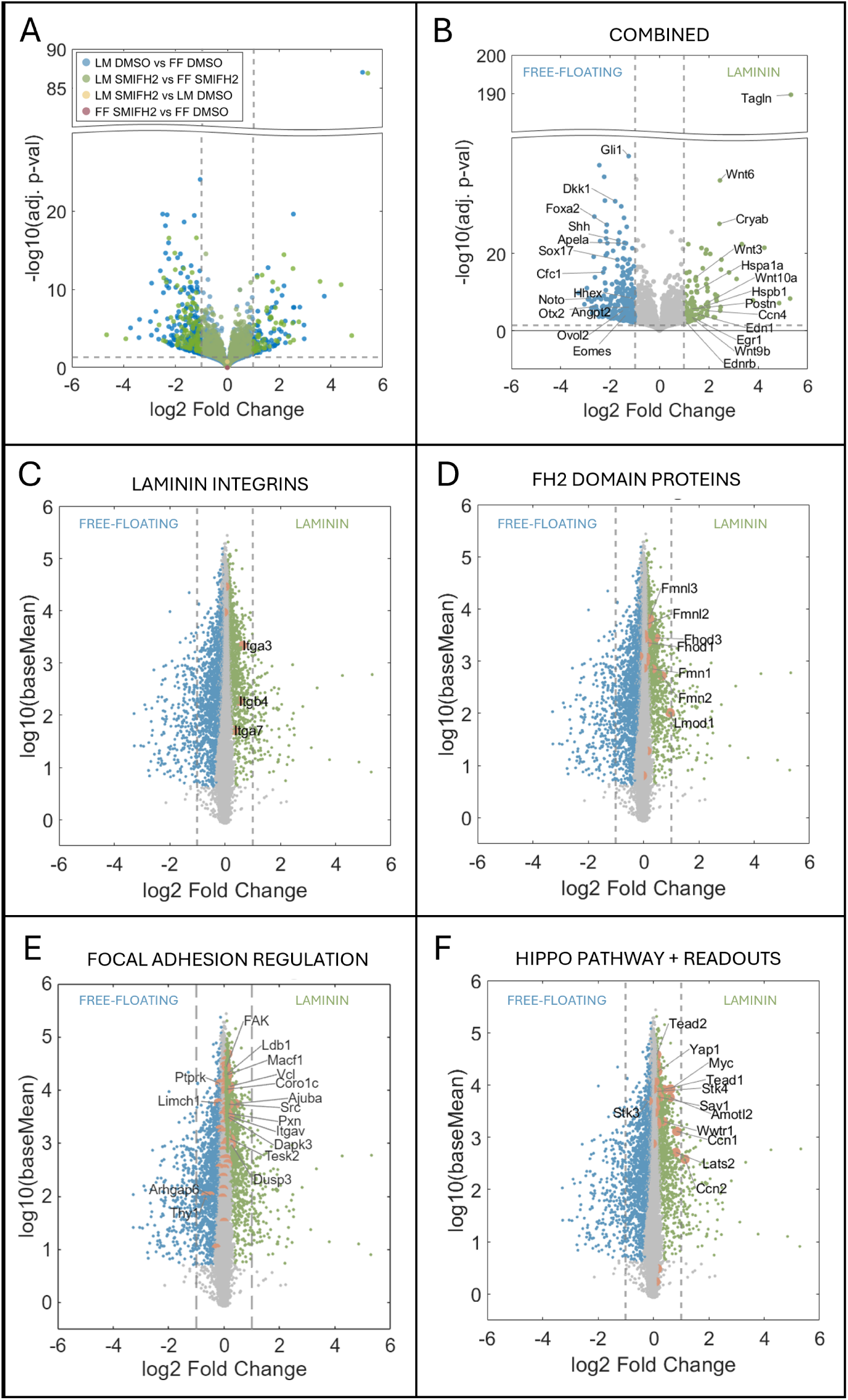
Laminin promotes the expression of genes involved in cell migration. A) Formin inhibition does not induce changes in gene expression. Volcano plot showing differentially expressed genes across conditions. Blue: laminin DMSO vs free-floating DMSO; green: laminin SMIFH2 vs free-floating SMIFH2; yellow: laminin SMIFH2 vs laminin DMSO; pink: free-floating SMIFH2 vs free-floating DMSO. Note the absence of differentially expressed genes in SMIFH2 comparisons (yellow and pink). Dashed lines indicate log2FC = ±1 and p-adj = 0.05. B) Volcano plot highlighting selected differentially expressed genes between laminin and free-floating conditions using the combined data (see methods). C-F) Plots highlighting expression levels of specific gene categories: laminin-binding integrins (C), FH2 domain-containing formins (D), focal adhesion regulators (E), and canonical YAP1 transcriptional targets (F). Coloured points indicate genes with p-adj < 0.05. Orange points in C-F highlight genes of interest within each category (only those with p-adj < 0.05 are labelled). Dashed lines indicate log2FC = ±1.

### Laminin biases cells toward migratory states while preserving axial identity

Next, we examined the gene expression changes associated with gastruloids growing on laminin. We observed changes in adhesion-related genes, including upregulation of the main laminin-binding integrins (Fig. 5C), several FH2 domain-containing formins including Fmn1, Fmn2, Fmnl2, and Fmnl3 (Fig. 5D), and focal adhesion (FA) regulators (Fig. 5E). Notably, while these expression changes are statistically significant, their magnitude is modest (generally with |log2FC|<1). These changes in the bulk transcriptomics could come from modest widespread changes, or by more substantial changes in small cell populations. Regardless, free-floating gastruloids express formins and extend filopodia (Fig. S9), yet achieve elongation through substrate-independent mechanisms that are insensitive to SMIFH2.

The transcriptional changes on laminin could be triggered by physical parameters associated with substrate adhesion, such as cell spreading. Consistent with this idea, we observe the upregulation of canonical YAP transcriptional targets (Ccn1, Ccn2, Myc; Fig. 5F), suggesting substrate-induced mechanotransduction consistent with previous reports linking cell spreading to YAP-dependent FA gene expression (Nardone et al., 2017).

Beyond adhesion and cytoskeletal changes, laminin also influenced developmental signalling. Gene Ontology analysis of the differentially expressed genes (|log2FC| > 1 and p-adj <0.05) shows changes in Wnt signalling (Fig. S12): Gastruloids on laminin showed upregulation of Wnt ligands and downregulation of Wnt antagonists, Nodal pathway components, and Shh targets (Fig. S13A-D). This transcriptional profile suggests laminin promotes a more posterior, mesenchymal identity consistent with the enhanced migratory behaviour observed in these conditions, while driving the downregulation of more anterior endodermal fates (Fig. S13E-F). However, Hox gene expression remained largely unchanged (Fig. S14), indicating that free-floating and laminin gastruloids acquire equivalent axial identities (posterior trunk) despite their distinct morphogenetic trajectories. Overall, the transcriptomic analysis shows that laminin biases cells toward migratory fates while the gastruloid preserves its axial identity. Critically, these cells achieve axis elongation through formin-dependent traction forces: a purely mechanical requirement that is dispensable in free-floating conditions, where elongation occurs through substrate-independent mechanisms.

### Focal adhesion-mediated traction drives axis elongation on laminin

Having established that formin-dependent filopodia are essential for cell stream formation specifically on laminin, we next investigated the role of FAs in this process. FAs serve as mechanical linkages between the ECM and the actin cytoskeleton, transmitting traction forces required for cell migration on substrates. Given the upregulation of FA regulators we observed in our transcriptomics analysis (Fig. 5E), we hypothesized that FA-mediated traction forces are specifically required for axis elongation on laminin.

We first examined the localization of phosphorylated FAK (pFAK), a marker of active focal adhesions, in gastruloids cultured on laminin. pFAK was enriched at the periphery of the gastruloid, within the elongating cell streams, and in cellular protrusions (Fig. 6A), consistent with active traction force generation in these regions.

**Fig. 6.**
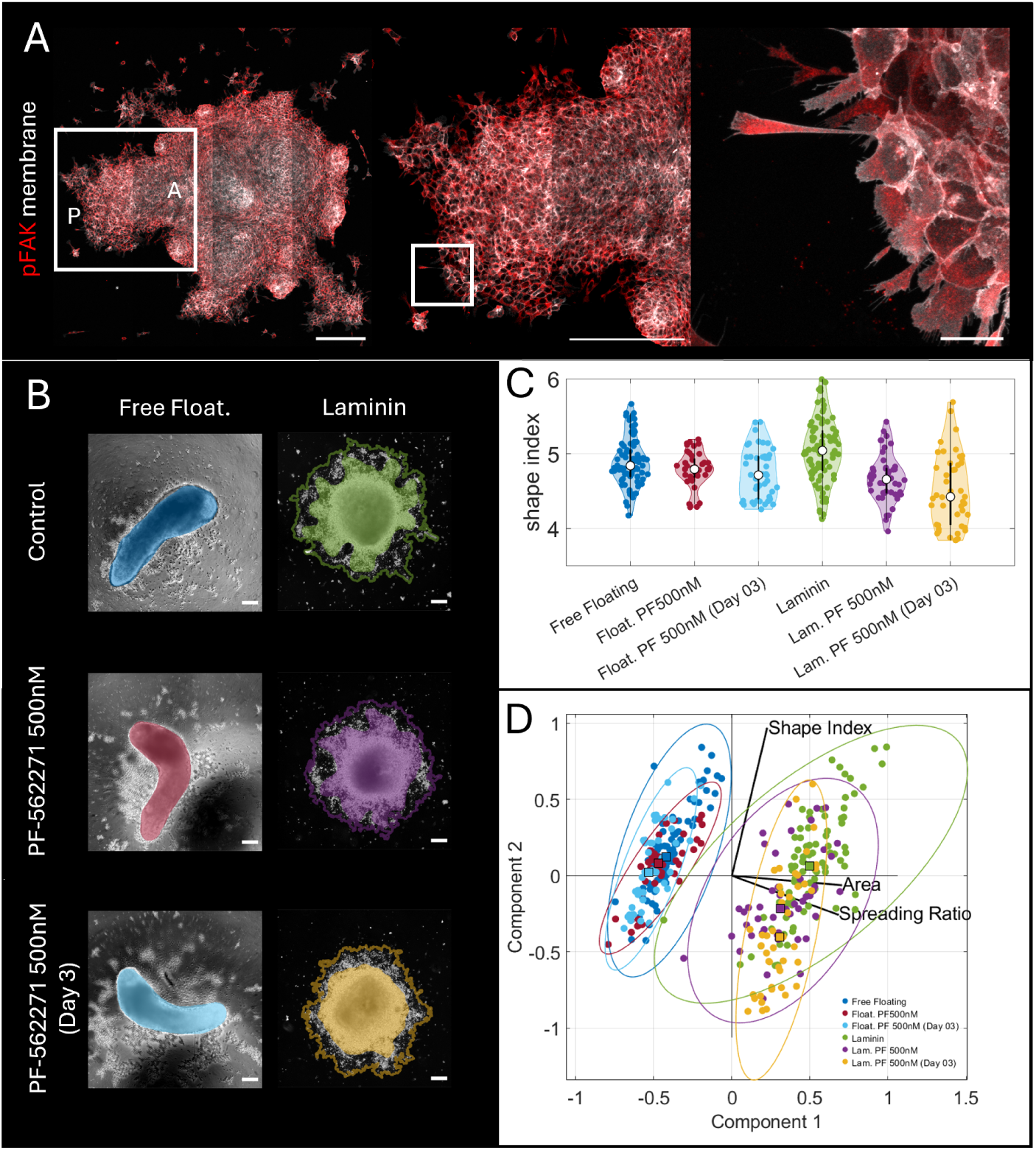
Focal adhesions are required for axis elongation only on laminin. A) Immunostaining for phosphorylated FAK (pFAK, red) and membrane marker (white) in gastruloids cultured on laminin at day 5. pFAK is enriched at the gastruloid periphery, within cell streams, and in cellular protrusions. Boxes indicate magnified regions. A and P indicate anterior and posterior poles, respectively. B) Representative images of freefloating and laminin gastruloids treated with DMSO (Control) or the FAK inhibitor PF-562271 (500 nM) added at day 4 or day 3.5. Coloured overlays indicate gastruloid outlines. C) Quantification of shape index across conditions. FAK inhibition reduces morphological complexity specifically on laminin, with earlier treatment (day 3) showing stronger effects. D) PCA. FAK inhibition shifts treated laminin gastruloids while free-floating gastruloids show no effect. See supplementary material for full statistical analysis. Colours indicate the same conditions across graphs. Scale bars: 200 *µm*. (A, left and center), 20 *µm*. (A, right), 200 *µm* (B).

**Fig. 7.**
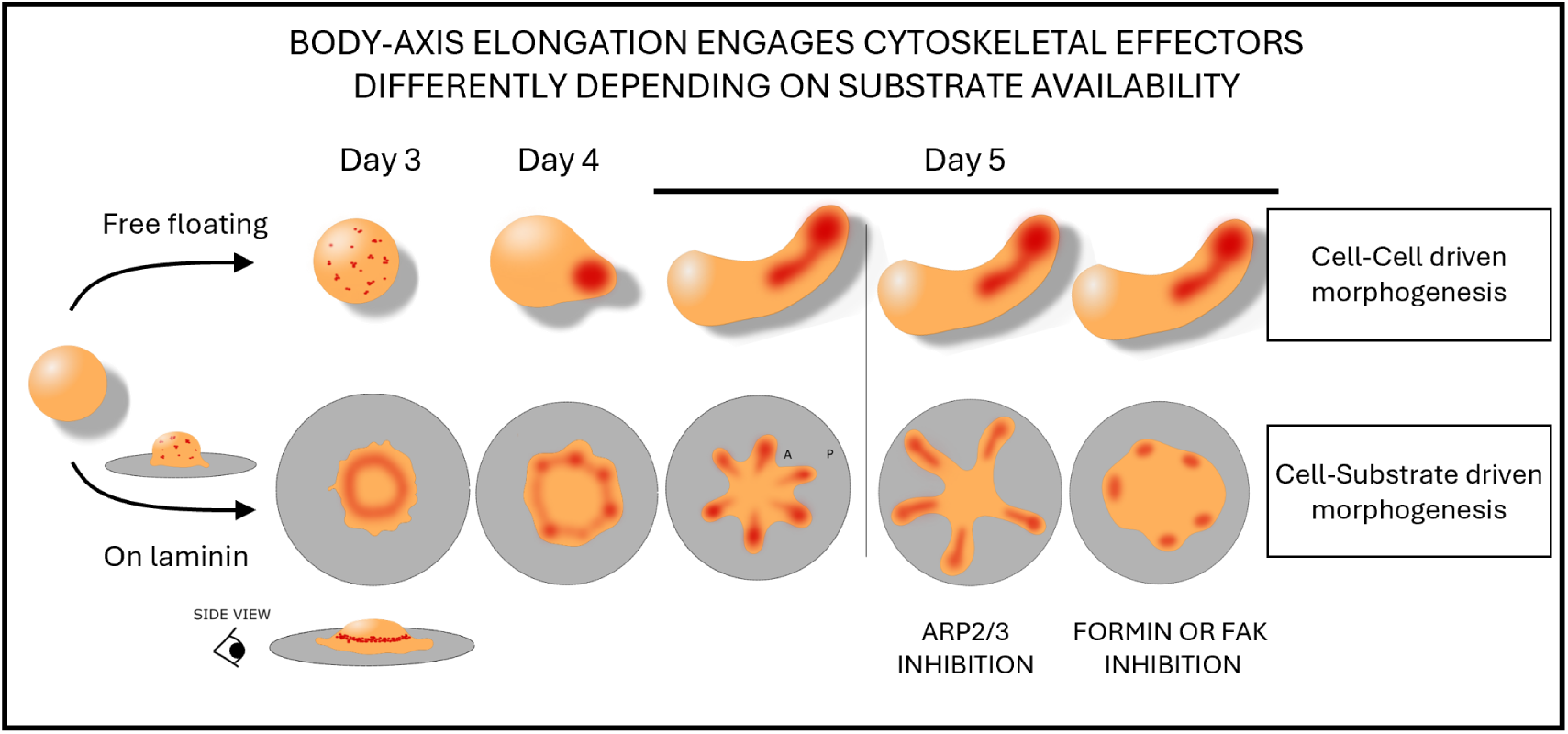
Adherent substrates reveal different morphogenetic routes to generate a posterior body axis in gastruloids. Summary of the findings: When grown on an adhesive substrate, gastruloids undergo body axis elongation using cell-substrate interactions regulated by Arp2/3, formins and focal adhesions. Instead, free-floating gastruloids elongate using cell-cell interactions that do not depend on ECM contact.

To functionally test the requirement for FAs, we treated gastruloids with PF-562271, a selective FAK inhibitor. FAK inhibition blocked body axis formation on laminin, with severity increasing with treatment duration, while free-floating gastruloids retained their normal elongated morphology (Fig. 6B-D).

These results suggest that body-axis elongation on laminin requires focal adhesion-mediated traction. Notably, FAK is highly expressed in both conditions (Fig. 5E, log2FC = 0.08), yet free-floating gastruloids do not need it for elongation. This suggests that morphogenetic plasticity could arise from the context-dependent engagement of pre-existing cytoskeletal components.

## Discussion

Our findings reveal that gastruloids possess a temporal cell-substrate adhesion profile that mirrors the sequential deposition of ECM components in the embryo. Gastruloids show strong affinity for laminin from the time of aggregation, independent of Chiron treatment, likely reflecting the primordial role of laminin in mammalian gastrulation: it is the first ECM protein to appear in mammalian embryos (Leivo et al., 1980; Cooper and MacQueen, 1983) and the only ECM component whose absence results in lethal phenotypes during gastrulation (Schéele et al., 2007). In contrast, the delayed, Chiron-dependent adhesion to fibronectin and collagens may reflect the requirement for mesoderm-associated EMT and subsequent ECM-specific integrin expression (Burdsal et al., 1993).

Early attachment to laminin constrains gastruloid morphogenesis, preventing the formation of a single body axis. While free-floating gastruloids consolidate mesoderm precursors into a single pole through extensive cell rearrangements (Hashmi et al., 2022; Gsell et al., 2025), gastruloids plated on laminin develop multiple body axes. This multipolar morphology likely arises because strong cell-substrate adhesion restricts cell mobility during days 2-4, trapping T/Bra-positive cells at their sites of induction around the gastruloid periphery. Unable to coalesce through collective rearrangements, each cluster of mesoderm precursors instead initiates local axis elongation, with each stream establishing proper anteroposterior patterning (Cyp26a1 at tips, Aldh1a2 in the body). Hox gene expression remains similar between conditions, indicating that both laminin and free-floating gastruloids acquire equivalent positional identity.

Our transcriptomics reveals upregulation of laminin-binding integrins, focal adhesion regulators, and canonical YAP transcriptional targets, consistent with substrate-induced mechanosensing (Nardone et al., 2017). We also observe enhanced Wnt signalling on laminin. This co-activation of YAP and Wnt pathways is consistent with evidence that ECM mechanical properties can modulate Wnt/*β*-catenin signalling through integrin-dependent mechanisms, and that YAP/TAZ can transcriptionally regulate Wnt pathway components (Astudillo, 2020). The resulting enhancement of Wnt signalling correlates with downregulation of anterior primitive streak derivatives, including definitive endoderm markers, and upregulation of mesenchymal markers, suggesting that mechanotransduction on laminin biases cells toward posterior migratory fates while maintaining trunk identity as marked by Hox expression. Finally, blocking tissue elongation on laminin through formin inhibition does not result in transcriptional changes, suggesting that the transcriptional programs driving cell fate specification are robust to disruption of morphogenesis (Bennabi et al., 2025).

By day 4, mesoderm cells have acquired the molecular machinery necessary for substrate-based migration, enabling the transformation of mesoderm clusters into actively elongating streams. This morphogenetic process shows specific cytoskeletal requirements: formin-dependent actin structures are essential for stream formation, while Arp2/3 activity paradoxically restricts elongation efficiency. Interestingly, similar cytoskeletal dependencies have been observed in cancer cell migration, where Arp2/3 inhibition increases linear migration velocity while formin inhibition blocks migration entirely (Monzo et al., 2016). In contrast, free-floating gastruloids are unaffected by inhibition of either formins or Arp2/3, indicating they employ different force-generation mechanisms: likely cell-cell interactions based on differential adhesion and cortical contractility (McNamara et al., 2024; Mayran et al., 2023; Fiuza et al., 2024; Oriola et al., 2024).

The substrate-specific phenotype of formin or FAK inhibition, in combination with the absence of transcriptional effects following formin inhibition, suggest that these proteins have a mechanical function that only manifests in substrate-based morphogenesis. Formins are necessary for the formation of filopodia and stress fibres, both actin structures with important roles in cell adhesion and migration (Fig. S15). Importantly, while ROCK inhibition, which blocks myosin 2-dependent stress fiber contractility (Totsukawa et al., 2000), impairs elongation in both free-floating and laminin conditions (Fig. S3), SMIFH2 selectively affects laminin-seeded gastruloids. This distinction suggests that the laminin-specific requirement involves formin-dependent actin nucleation rather than myosin 2-driven contractility. However, we note that SMIFH2 also inhibits some myosins (Nishimura et al., 2021), including myosin 10, which localises to filopodia tips, so the precise contribution of filopodia versus other formin-dependent structures remains to be determined.

Nevertheless, focal adhesions can form at filopodia tips during collective cell migration (Bischoff et al., 2021), providing mechanical anchorage for force transmission. Furthermore, FAK-null mice can implant and initiate gastrulation; however, body-axis elongation was dramatically retarded by E8.5 (Furuta et al., 1995). FAK deficiency did not impair the differentiation potential of mesoderm cells or their proliferation, but produced a drastic reduction in their migratory speed and directionality (llić et al., 1995). Notably, FAK-deficient cells exhibited an increased number of focal adhesions rather than fewer, indicating that FAK promotes focal adhesion turnover rather than assembly (llić et al., 1995). This is consistent with the failure of cell stream formation we observe upon FAK inhibition on laminin, where cells remain attached to the substrate but fail to elongate. Together, these results suggest that substrate-based mesoderm migration contributes to body-axis elongation in the embryo.

Notably, T/Bra-positive cells within free-floating gastruloids generate active protrusions dependent on non-canonical Wnt signaling (Anlaş et al., 2024; Fiuza et al., 2024). Blocking non-canonical Wnt prevents coalescence of T/Bra clusters (Fiuza et al., 2024), suggesting this pathway drives the mechanical changes underlying mesoderm progenitor sorting. Together with our findings, this might suggests that non-canonical Wnt induces an excitable cytoskeleton that generates protrusions and alters mechanical properties, driving tissue elongation through cell-cell interactions in free-floating conditions or cell-substrate interactions when adherent to ECM.

Although the plastic properties of vertebrate gastrulation can be studied in the embryo (Chuai et al., 2023; Serra et al., 2023; Serrano Nájera et al., 2024), embryonic development is a tightly regulated, robust process that may mask the latent morphogenetic capacities of embryonic tissues (Serrano Nájera and Weijer, 2023). For these reasons, in vitro systems, where embryonic constraints are relaxed, may be better suited to reveal this hidden potential. For instance, cerebral organoids generate ventricle-like lumens through cavitation (Lancaster et al., 2013), whereas in vivo these structures derive from the neural tube lumen, which forms through folding morphogenesis. Yet when appropriate geometrical and adhesive constraints are applied, folding-based neurulation can be recapitulated (Karzbrun et al., 2021). Analogously, mouse embryonic stem cell aggregates normally form lumens through hollowing in an E-cadh-dependent process, but in the absence of E-cadh they generate lumen-like cavities via a closure mechanism instead (Liang et al., 2022). However, these studies identified alternative morphogenetic outcomes without systematically comparing the cellular requirements of each mode. Here, we first show that gastruloid elongation can proceed through alternative morphogenetic modes, and by applying the same perturbations across both conditions, we demonstrate that each mode has distinct cytoskeletal dependencies.

Together with these examples, our findings demonstrate that cells retain the capacity for multiple morphogenetic programs to achieve structurally similar outcomes, perhaps through a process reminiscent of phenotypic accommodation (West-Eberhard, 2005). This flexibility may have evolutionary significance: it challenges deterministic views of morphogenesis and suggests that evolutionary innovations may arise not from new gene-regulatory programs, but from environmentally induced shifts in how existing cytoskeletal machinery is used, revealing latent mechanical capabilities without requiring genetic novelty. Understanding that morphogenetic outcomes can be plastically redirected by tuning mechanical context suggests new strategies for tissue engineering through the control of physical properties and cytoskeletal activity.

In conclusion, our work demonstrates that gastruloids can achieve body-axis elongation through different morphogenetic strategies depending on whether or not cells engage substrates to generate force. This shows that similar developmental outcomes can emerge from distinct force-generation strategies, providing potential raw material for evolutionary diversification. On the practical side, these findings suggest that desired morphologies might be achieved by triggering latent morphogenetic programs.

## Acknowledgments

GSN acknowledges support from Leverhulme Trust Early Career Fellowship (ECF-2022-474). AD and BJS acknowledges support from the a Wellcome Trust Discovery Award (225360/Z/22/Z). We thank the Steventon lab members, Marta Urbanska for their support and enlightening discussion, Imen Lassadi and Oskar Batty for sharing of the SBN line and Oluwaseun Ogundele for her support with the processing transcriptomics data.

## Author contributions

CRediT taxonomy terms for the individual contributions.

GSN: Conceptualisation, Software, Validation, Formal Analysis, Investigation, Data Curation, Writing – Original Draft, Writing – Review & Editing, Visualisation, Supervision, Funding acquisition. AD: Validation, Formal Analysis, Investigation, Data Curation, Writing – Original Draft, Visualisation. BJS: Conceptualisation, Resources, Project administration, Writing – Review & Editing, Funding acquisition.

## Supplementary material

### Materials and methods

All materials used are listed [here](https://www.dropbox.com/scl/fi/mbfp7xwrlz287u9nzx66y/reagents_list_30062025.xlsx?rlkey=wdkz9uevf7w61kat7rqy9xa6s&st=6et8wlsw&dl=0).

### Cell culture

mESC were cultured at 37°C, 5% CO2 in 2i LIF (N2B27 (NeuroBasal medium, DMEM F-12 no glutamine, N2 —homemade (Mulas et al., 2019)—, B27, 2-Mercaptoethanol), l-glutamine, 3µM CHIR99021, 1µM PD0325901 and 0.1µg ml-1 LIF). All lines were cultured in filter-capped tissue culture flasks coated with 0.1% gelatin. Cell culture media was changed everyday, and cells were passaged every 2 days. Cells were routinely tested for mycoplasma. See supplementary table X for compounds references.

### Cell lines

The lines used in this study are the parental ES-E14TG2a, the triple reporter ES-E14TG2a containing Sox1-P2A-eGFP and Bra-P2A-mCherry (Deluz et al., 2016) modified with H2B:miRFP670 (SBN), and ES-E14TG2a with CAG::GPI-GFP (GPI-GFP) (Rhee et al., 2006).

### Gastruloid culture

Gastruloids were generated following a previously published protocol (Baillie-Johnson et al., 2015). A total of 200–300 mouse embryonic stem cells (mESCs) were aggregated in 40 *µL* of N2B27 medium supplemented with Penicillin-Streptomycin (1%) and L-glutamine (referred to as supplemented N2B27) using a cell-repellent, U-bottom 96-well plate (650970, Greiner). Cells were allowed to aggregate for 48 hours, after which 150 *µL* of supplemented N2B27 containing 3 *µM* CHIR99021 (CHIRON pulse) was added to each well. Gastruloids were incubated in this medium for 24 ± 2 hours, after which 150 *µL* of medium was replaced daily with fresh supplemented N2B27. Gastruloids were maintained in culture for a total of 5 days. For the activin A experiment, the chiron pulse from the original protocol was replaced by an Activin A (100ng/mL) pulse of 24h, from 48h to 72h.

### 2D gastruloid protocol

Gastruloids were generated as described above. On day 2, 35 mm glass-bottom dishes were coated with 14 *µg/mL* Laminin 111 (L2020, Sigma) for 1 hour at 37 °C. Laminin solution is prepared by diluting the stock solution in PBS without calcium and magnesium (PBS -/-) to a final concentration of 14 *µg/mL*. Gastruloids were collected using a cut 200 *µL* pipette tip and allowed to settle at the bottom of an Eppendorf tube for a few seconds. The medium was then carefully removed, and gastruloids were resuspended in supplemented N2B27 medium containing 3 *µM* CHIR99021. They were subsequently transferred, using a cut tip, into the Laminin-coated dish containing fresh medium. To prevent fusion, individual gastruloids were gently separated from one another using a cut tip. The dish was then placed in the incubator, which remained closed for at least one hour to allow for proper attachment without disturbance. After 24 hours, gastruloids typically appeared flatter. From this point, medium was changed daily as described above until day 5. In the same manner, fibronectin coating is realized by diluting Fibronectin stock solution in PBS-/- to a final concentration of 12.5µg/mL. The solution is transferred to a glass-bottom dish (400µL for a 20mm dish) and incubated for 1 hour at 37°C. Before seeding, the solution is removed from the dish and washed once with PBS-/-. For collagens coatings, solutions are kept on ice until use. Collagen I is prepared by diluting the stock solution in 0.02N acetic acid to a final concentration of 12.5µg/mL overnight at 4 °C. The solution is then added to the dish and left to incubate at RT for one hour. Dishes are rinsed twice with PBS to remove acid prior gastruloids plating. Collagen IV is prepared by diluting the stock solution in cold deionized water to a working concentration of 12.5µg/mL. The subsequent steps are identical to those described for Collagen I.

### Mosaic generation

Mosaics were generated following the 3D gastruloid protocol described above but using a cell suspension composed of 95% E14 and 5% GPI-GFP cells.

### Chemical perturbations

Inhibitor treatments were carried out between 84h and 96h. CK666 (SML0006-5MG, Sigma), is resuspended in DMSO at a 10mM concentration upon reception and stored at 4°C. SMIFH2 (HY-16931, MedChemExpress), is resuspended in DMSO at a 10mM concentration upon reception and stored at -80°C. GSK269962A (S7687-SEL, Selleckchem), is resuspended in DMSO at a 10mM concentration upon reception and stored at -80°C. Inhibitor concentrations are as follows: CK666 (10µM and 50µM), SMIFH2 (10 and 25µM), GSK269962A (0.1µM).

### Immunofluorescence staining

Gastruloids for staining are transferred to an Eppendorf using a cut tip, they are then washed once with PBS-/- before being fixed with 4% PFA overnight at 4°C (or 3 hours at room temperature). The 2D gastruloid staining protocol is identical to the 3D protocol, except that all the steps are performed directly in the dish. On the following day, gastruloids are washed twice with PBS-/-, prewashed with PBSFT ( PBS -/- with 10%FBS (F9665-500ML, ThermoFisher) and 0.2% Triton X-100), washed three times with PBSFT at room temperature (10 minutes), and incubated in PBSFT for 1h at 4°C on an orbital shaker for blocking. Primary antibody solutions are prepared in PBSFT and used to replace the blocking solution once blocking is complete. Gastruloids are incubated overnight at 4°C on an orbital shaker. For 2D gastruloids, to avoid wasting antibodies, the full dish is not filled with solution. Instead, a dome of antibody solution covering the glass part of the dish is sufficient (around 400µL for a 20mm diameter glass-bottom dish). Next day, the primary antibody solution is washed as follows: 2×5 min in PBSFT (RT), 3 x 10 min PBSFT (4°C, orbital shaker), 4x 30 min (4°C, orbital shaker). Secondary solutions are prepared in PBSFT with nuclear (Sytox / DAPI) or / and Phalloidin. PBSFT wash is replaced with secondary solutions and the tubes are covered in foil and incubated overnight at 4°C on an orbital shaker. Next day, secondary solutions are washed as described for the primary solutions. 3D gastruloids are placed in clearing solution —Scale S4,(Hama et al., 2015)— for 24 hours before mounting. 2D gastruloids are directly imaged or stored in PBS and sodium azide (0.05%) until imaging. A list of the antibodies used in this study can be found in the supplementary materials here.

### Hybridization Chain Reaction

Fixed gastruloids are dehydrated in methanol as follows: 5 minutes per wash at room temperature: 25% MetOH / 75% PBST (PBS -/-, 0.2% Triton X-100), 50% MetOH / 50% PBST, 75% MetOH /25% PBST, 100% MetOh twice. Gastruloids are then incubated overnight at -20°C. Next day, probe hybridization buffer is pre-warmed in the incubator and amplification buffer is brought at room temperature. Gastruloids are rehydrated with a serie of 5 minutes washes, 75% MetOH /25% PBST, 50/50, etc. Once the last wash is removed, gastruloids are pre-hybridized using warm probe hybridization buffer for 30min at 37°C. Probe solution is prepared by adding 2pmol of each probe set into hybridization buffer at 37°C, the solution is transferred to the gastruloids and incubated overnight at 37°C. Next day, pre-warm probe wash buffer in the incubator and wash the gastruloids 4 x 15min at 37°C. Wash 2 x 5 min with 5 X SSCT at room temperature. Amplification buffer is added to the gastruloids to pre-amplify ( 5 min, RT). Prepare hairpin solutions following supplier’s recommendation and add the solution to the gastruloids, incubate overnight in the dark at room temperature. The next day, remove excess hairpins by washing 3×15 min with SCCT at room temperature. Samples can be stored at 4°C until imaging / DAPI staining overnight in PBST. All HCR probes were bought from Sigma Aldrich.

### Confocal microscopy

Individual gastruloids were imaged using a Zeiss LSM700 confocal microscope, typically with a 20X objective, 0.5× zoom, and 2 *µm* axial slicing. For *in vivo* experiments, we used a humidified incubation chamber (5% CO2, 37°C) with a time step of 5 min for 12 h and an interval between slices of 2µm.

### High-throughput microscopy

Individual gastruloids were imaged using a Nikon Eclipse Ti-E inverted bright-field microscope equipped with a motorized stage, an Orca Flash 4.0 camera, a Lumencor SPECTRA X Light Engine with internal 391/22 nm, 475/28 nm, 555/28 nm, and 635/22nm excitation filters and a humidified incubation chamber (5% CO2, 37 °C). To locate each sample, we performed tile scans of individual positions on a 96-well plate (2×2) or 35 mm glass-bottom dish (7×7) at 5X magnification. Gastruloid positions were then selected using our custom MATLAB-based tool, which enables loading of multi-point microscopy data and identification of gastruloids either manually through user selection or via optional AI-assisted detection powered by Cellpose segmentation, all through an intuitive graphical interface. Once individual gastruloid positions had been recorded, we acquired single snapshots or time-lapse images with a 20X objective, capturing phase contrast and fluorescence images with appropriate illumination channels as needed.

### High-throughput image analysis

We developed a MATLAB-based computational pipeline that integrates multiple tools for the segmentation and analysis of gastruloids. We used Cellpose models (Pachitariu and Stringer, 2022) trained from scratch to detect the variable morphologies of gastruloids cultured in free-floating conditions or on ECM-coated dishes. To identify the mesenchymal migration front, we utilized probability maps from a pretrained Cellpose model (CP) to detect individual cells and extracted areas with probability > 0.35. For quantitative parameter extraction (area, elongation, circularity, fluorescence intensity, etc.), as well as the LOCO-EFA descriptors (Sánchez-Corrales et al., 2018), we fed the Cellpose segmentations and original images into MOrgAna (Gritti et al., 2021). Finally, diagnostic visualization, data representation, and statistical analysis were performed using MATLAB. Code for this pipeline can be accessed here.

### Particle Image Velocimetry

Velocity fields were computed from confocal time-lapse movies (time frame = 5 min) using PIVLab v2.56 for MATLAB (Thielicke and Sonntag, 2021) with two passes of 64×64 and 32×32 pixel interrogation windows with 50% overlap with default pre- and post-processing parameters.

### Lamellipodia and Filopodia quantification

Lamellipodia were manually delineated using the polygon tool in FIJI (Schindelin et al., 2012). Individual polygon coordinates were exported using a custom ImageJ macro, and the area of each polygon (lamellipodia) was computed and visualised using MATLAB. Filopodia were measured through a similar procedure, using the free-line tool in FIJI to manually trace them.

### Cell division angle quantification

Division angles between daughter cell pairs were quantified at the moment of cytokinesis using confocal time-lapse movies acquired between days 4 and 5 of gastruloid development on laminin-coated substrates. The quantification procedure involved the following steps:

1. Daughter cell pair coordinates were manually identified using the multipoint tool in FIJI, and the resulting coordinates were exported to MATLAB for analysis.
2. The main elongation axis of each cell stream was defined by drawing a line along the stream’s length on the final frame of the time-lapse movie using MATLAB’s drawline function.
3. Cell divisions were filtered to include only those occurring within the “neck” region of the cell stream, which was defined by manual delineation using MATLAB’s drawpolygon function.
4. The angle between each cell division axis and the main elongation axis of the corresponding cell stream was calculated for all samples and experimental conditions.

### Statistical Analysis

All results were compiled into matrices showing adjusted p-values for all pairwise comparisons and exported as Excel files. See the Supplementary Material to find the statistical analysis associated with each figure.

### Analysis for obtained from the high-throughput analysis pipeline

Statistical analyses were performed using MATLAB using a non-parametric approach. The Wilcoxon rank-sum test (Mann-Whitney U test) was conducted between all possible group combinations. This non-parametric test ranks all observations from both groups being compared and tests whether the distributions differ significantly, making it suitable for data that may not meet the assumptions of parametric tests. To account for multiple comparisons in the pairwise testing phase, p-values were adjusted using the Bonferroni correction method, where each p-value was multiplied by the total number of pairwise comparisons and capped at 1.

### Analysis of lamellipodia and filopodia

MATLAB’s implementation of the Wilcoxon rank-sum test was conducted between all possible group combinations. p-values were adjusted using the Bonferroni correction method.

### Analysis for cell division angle data

Cell division angle analysis was performed using circular statistics appropriate for axial data using the Circular Statistics Toolbox (Berens, 2009) for MATLAB. Division angles were measured as the angle between the division axis and a reference orientation, where angles *θ* and *θ* + *π*represent the same division axis. To apply circular statistical methods to axial data, all angles were converted to circular representation by doubling (*θ →* 2*θ*), transforming the axial period of *π* to a circular period of 2*π*. This transformation preserves the axial symmetry while enabling the use of standard circular statistical functions. To test the alignment of the distribution with a preferred direction, the V-test (Rayleigh test for a specified direction) was used to test for significant concentration around specific orientations. This non-parametric test evaluates the null hypothesis that data are uniformly distributed against the alternative hypothesis of concentration around a specified direction. We tested both groups for alignment at 0 (horizontal divisions) and *π/*2 (vertical divisions) orientations with respect to the elongation axis.

### Computational Surface Extraction

Active myosin resides in the most apical compartment of the cells. For this reason, maximum projections may not accurately represent the apical myosin, including the myosin cables. To tackle this problem we computationally acquire a 2D surface of the most apical side of the sample. To extract the apical surface of the embryo from the confocal image volumes, we used a custom MATLAB script based on the square gradient focusing algorithm (Eskicioglu and Fisher, 1995; Serrano Nájera, 2021; Serrano Nájera et al., 2024). The steps are as follows:

1. Tile the confocal volume into columns.
2. Apply the square gradient focusing algorithm to each column to find the surface position.
3. Generate a height map by finding the depth of the fastest change in the image sharpness (peak of the second derivative) for each column.
4. Apply a smoothing filter to the height map.
5. Use the smoothed height map to section the image volume and produce a 2D image of the tissue surface.

### Transcriptomics analysis

#### Experimental design and sample collection

RNA-sequencing was performed on gastruloids cultured under four conditions: free-floating with DMSO (FF-DMSO), free-floating with SMIFH2 25 µM (FF-SMIFH2), laminin with DMSO (LM-DMSO), and laminin with SMIFH2 25 µM (LM-SMIFH2). For each condition, three biological replicates were collected at day 5 (120 h), with each replicate consisting of pooled gastruloids (n = 24 per replicate for free-floating, n = 20 per replicate for laminin). SMIFH2 or DMSO vehicle was added at 84 h. Gastruloids were collected, washed in PBS, and lysed in TRIzol. Total RNA was extracted using phenol-chloroform phase separation followed by isopropanol precipitation.

#### Library preparation and sequencing

RNA-sequencing libraries were prepared and sequenced at the Cambridge Stem Cell Institute sequencing facilities. All 12 samples were processed in a single batch. Starting from 300 ng of total RNA per sample, poly(A) mRNA was isolated using the NEB Poly-A Magnetic Isolation Module, which employs oligo d(T) beads to capture polyadenylated transcripts. External RNA controls (ERCC spike-ins) were included prior to library preparation. Strand-specific libraries were generated using the NEBNext Ultra II Directional RNA Library Prep Kit. Libraries were quantified, combined in an equimolar pool, and sequenced on an Illumina NovaSeq X platform using a paired-end 150 bp (PE150) configuration. Sequencing was performed to an average depth of approximately 30–50 million paired-end reads per sample. Base calling and demultiplexing were carried out by the sequencing facility prior to downstream bioinformatic analysis.

#### Bioinformatic processing

Raw sequencing data were converted to FASTQ format using Illumina bcl2fastq software. Sequencing quality was assessed using FastQC, and quality summaries were compiled using MultiQC (Ewels et al., 2016). Adapter trimming and quality filtering were performed using Trim Galore (v0.6.1) in paired-end mode, retaining reads with a minimum length of 25 bp (Martin, 2011). Post-trimming read quality was confirmed using FastQC.Trimmed reads were aligned to the mouse reference genome (mm10 / GRCm38) using BWA-MEM (Li and Durbin, 2009), with the -M flag enabled. SAM files were converted to BAM format and processed using SAMtools v1.13 (Li et al., 2009). Mate information was added using samtools fixmate, followed by coordinate sorting. PCR duplicates were identified and removed using samtools markdup -r, generating duplicate-filtered BAM files. Reads mapping to the mitochondrial chromosome (chrM) were excluded prior to downstream analysis. BAM files were indexed, and alignment quality was assessed using samtools flagstat at multiple processing stages (Li et al., 2009). Gene-level quantification was performed using featureCounts, assigning uniquely mapped reads to annotated exons using the Ensembl Mus musculus GRCm38 gene annotation (Liao et al., 2014).

#### Differential expression analysis

Differential gene expression analysis was carried out in R using DESeq2 (Love et al., 2014). Raw counts were normalised using the median-of-ratios method, and lowly expressed genes were filtered prior to model fitting. To assess the transcriptional effects of both culture substrate and SMIFH2 treatment, four pairwise comparisons were initially performed using three biological replicates per condition: (1) LM-DMSO vs FF-DMSO, (2) LM-SMIFH2 vs FF-SMIFH2, (3) FF-SMIFH2 vs FF-DMSO, and (4) LM-SMIFH2 vs LM-DMSO. P-values were adjusted for multiple testing using the Benjamini–Hochberg method, and genes with an adjusted p-value below 0.05 were considered significantly differentially expressed. Log2 fold-change shrinkage was applied using DESeq2 shrinkage estimators for visualisation and ranking.

Comparison of DMSO- and SMIFH2-treated samples revealed no differentially expressed genes in either free-floating or laminin conditions (Fig. 5A, Fig. S11), confirming that SMIFH2 treatment does not alter the transcriptional state of gastruloids (Sup. Table 1). Given this transcriptional equivalence, and that all samples were processed in the same sequencing batch, DMSO- and SMIFH2-treated samples were pooled within each culture condition to increase statistical power for detecting substrate-dependent gene expression changes. This resulted in six biological replicates per condition (free-floating vs laminin). The pooled analysis yielded substantially improved statistical significance (Fig. 5B, Sup. Table 1), confirming that DMSO- and SMIFH2-treated samples are not merely lacking detectable differences but are transcriptionally consistent within each culture condition.

#### Gene Ontology Analysis

Gene Ontology enrichment analysis was performed on genes with |log2FC| > 1 and adjusted p-adj < 0.05 using GeneCodis 4.0 (Garcia-Moreno et al., 2022).

#### Gene category annotations

To examine expression changes in specific functional categories, we curated gene lists from Gene Ontology (GO) and published single-cell RNA-sequencing datasets.

##### Gene category annotations

To examine expression changes in specific functional categories, we curated gene lists from Gene Ontology (GO), published datasets, and manual literature curation.

##### GO-derived categories

Genes associated with focal adhesion regulation, filopodium assembly, and stress fiber assembly were obtained by combining related GO Biological Process terms. For filopodium assembly, we merged genes annotated with the following terms: ‘filopodium assembly’ (GO:0046847), ‘regulation of filopodium assembly’ (GO:0051489), ‘positive regulation of filopodium assembly’ (GO:0051491), and ‘negative regulation of filopodium assembly’ (GO:0051490). Similar approaches were used for focal adhesion regulation and stress fiber assembly, combining parent terms with their positive and negative regulatory sub-terms. GO annotations were obtained from the org.Mm.eg.db R package.

##### Cell type markers

Gut endoderm and mesenchyme marker genes were derived from the single-cell molecular atlas of mouse gastrulation (Pijuan-Sala et al., 2019). Differentially expressed genes defining gut endoderm and mesenchyme clusters at embryonic day 8.5 were extracted from the published dataset.

##### Other categories

Laminin-binding integrins, FH2 domain-containing formins, YAP transcriptional targets, and signalling pathway components (Wnt ligands, Wnt secreted antagonists, Nodal core signature, Shh targets) were manually curated from published literature.

### Manuscript preparation

During the preparation of this work the authors used Claude Sonnet 4.0 in order to check grammar and readability. While using this tool/service, the authors reviewed and edited the content as needed and take full responsibility for the content of the publication.

## Supplementary movies

- Movie 1 Gastruloid on laminin from day 4 to 5. Scale bar 500 *µm*.
- Movie 2 Gastruloid on laminin from day 4 to 5 in the presence of CK666 50 *µM*. Scale bar 500 *µm*.
- Movie 3 Gastruloids on laminin from day 4 to 5 with CK666 at 25 and 50 *µM*. Scale bar 500 *µm*.
- Movie 4 Gastruloid on laminin from day 4 to 5 in the presence of SMIFH2 25 *µM*. Scale bar 500 *µm*.

## Supplementary Tables

**Supplementary Table 1.** Differentially expressed genes across conditions. Genes with adjusted p-adj *<* 0.05 and |log2FC*| >* 1 were considered differentially expressed.

| Comparison | Replicates | Upregulated | Downregulated |
| --- | --- | --- | --- |
| *Effect of SMIFH2 treatment* |  |  |  |
| Free-floating: SMIFH2 vs DMSO | 3 vs 3 | 0 | 0 |
| Laminin: SMIFH2 vs DMSO | 3 vs 3 | 0 | 0 |
| *Effect of substrate* |  |  |  |
| Laminin vs Free-floating (DMSO only) | 3 vs 3 | 74 | 190 |
| Laminin vs Free-floating (SMIFH2 only) | 3 vs 3 | 55 | 181 |
| Laminin vs Free-floating (pooled) | 6 vs 6 | 116 | 266 |

## Supplementary Figures

**Supplementary Fig. 1.**
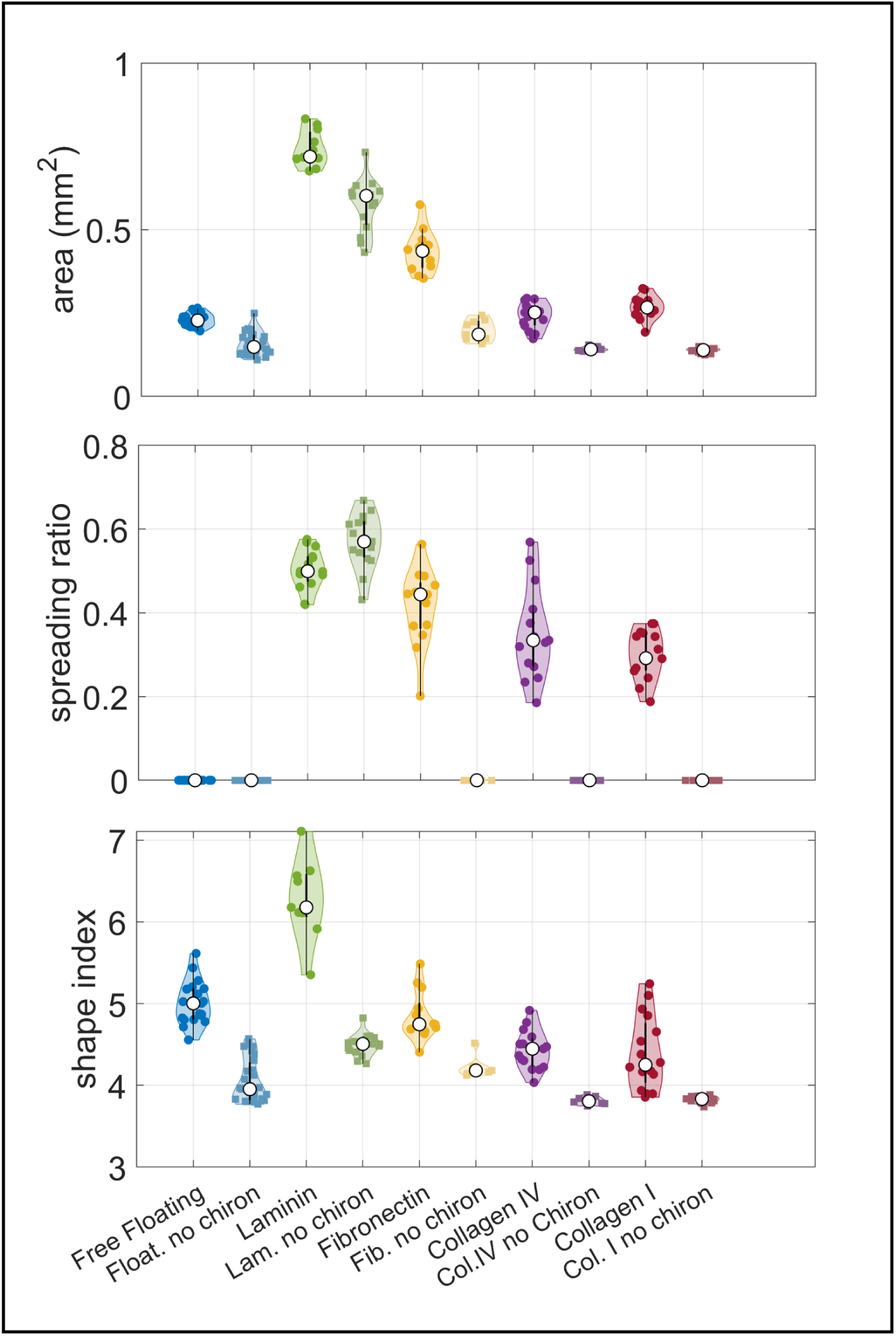
Characterisation of gastruloids grown on different substrates. Analysis of area, migratory abilities (spreading ratio) and elongation (shape index) on day 5 gastruloids grown on different coatings in the presence/absence of Chiron. See the supplementary material for the full statistical analysis.

**Supplementary Fig. 2.**
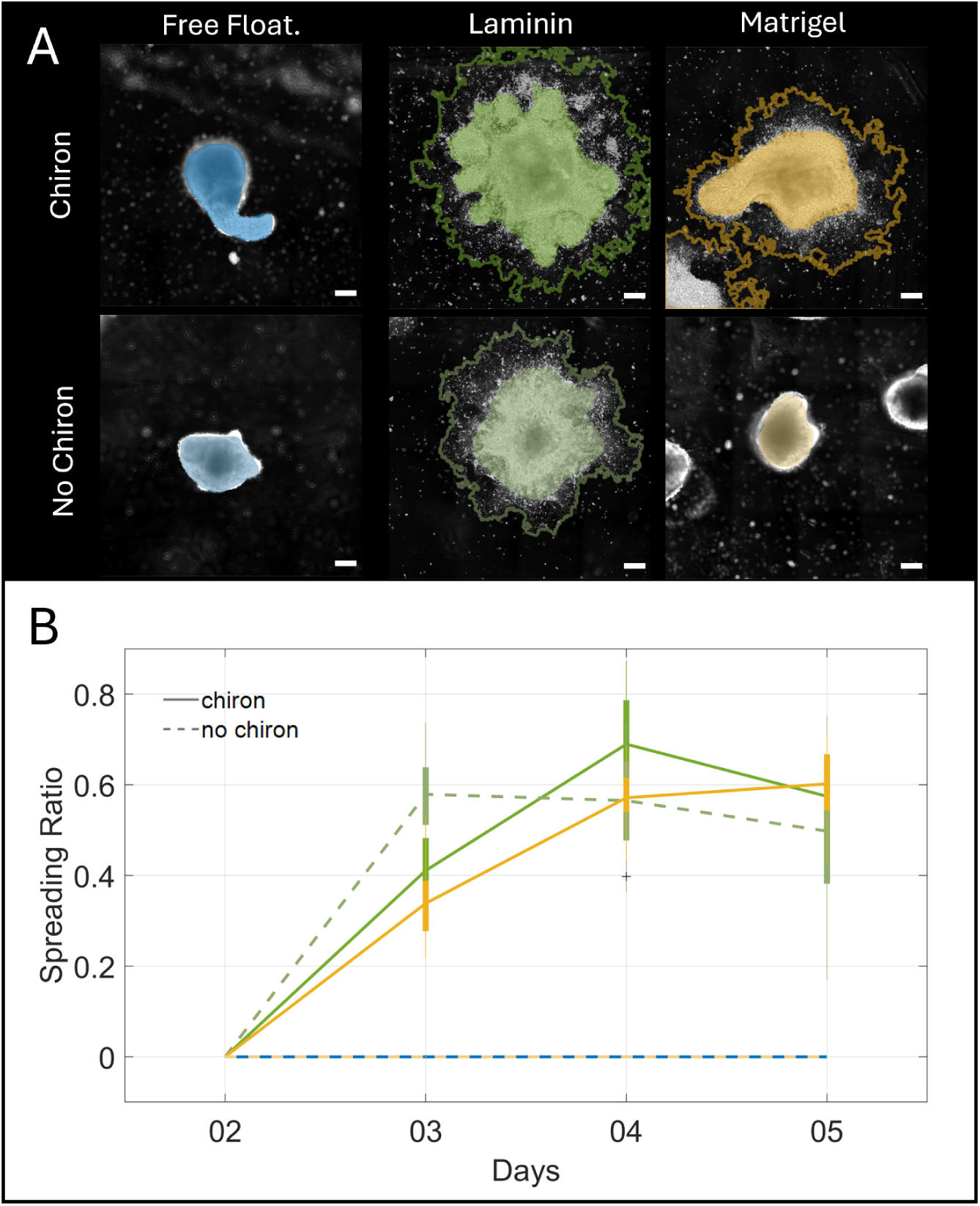
Adhesion to Matrigel requires chiron. A) Representative morphologies depending on the substrate and the presence of chiron. B) Analysis of the spreading capability of gastruloids on different substrates from day 2 to 5. Scale bars 200 *µm*.

**Supplementary Fig. 3.**
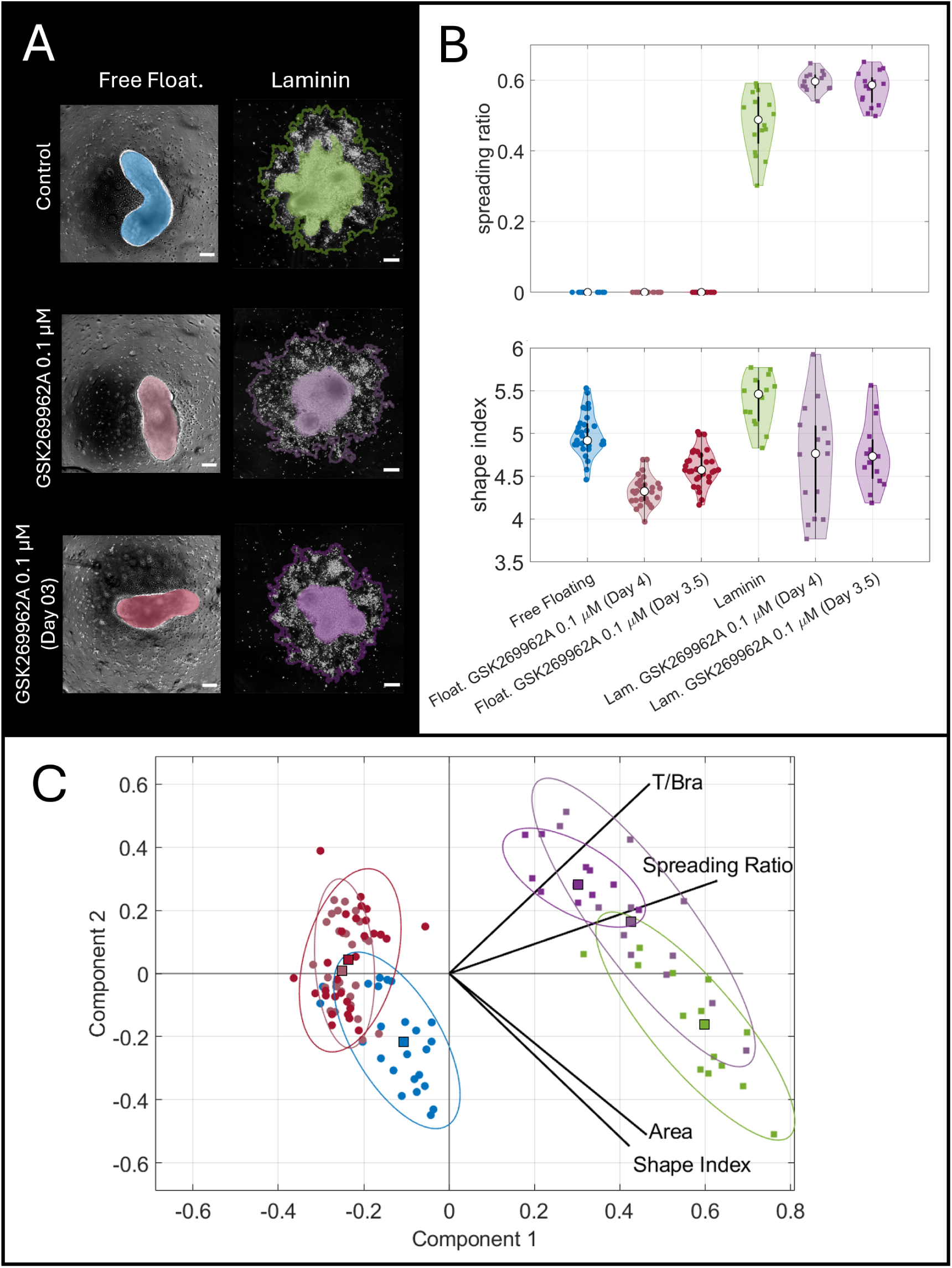
Myosin contractility is necessary for elongation in free-floating and laminin gastruloids. A) Gastruloid morphologies on different substrates with or without the ROCK inhibitor GSK269962A 0.1 *µM* added on day 3 or 4. B) Top: Analysis of the spreading capabilities after ROCK inhibition. Cell migration is enhanced on laminin. Bottom: Quantification of the shape index after ROCK inhibition. Free-floating and laminin gastruloids show a reduced elongation. C) Principal Component Analysis integrating different properties on day 5. Note that treated free-floating gastruloids do not overlap with the controls. Colours indicate the same conditions across graphs. See the supplementary material for the full statistical analysis.

**Supplementary Fig. 4.**
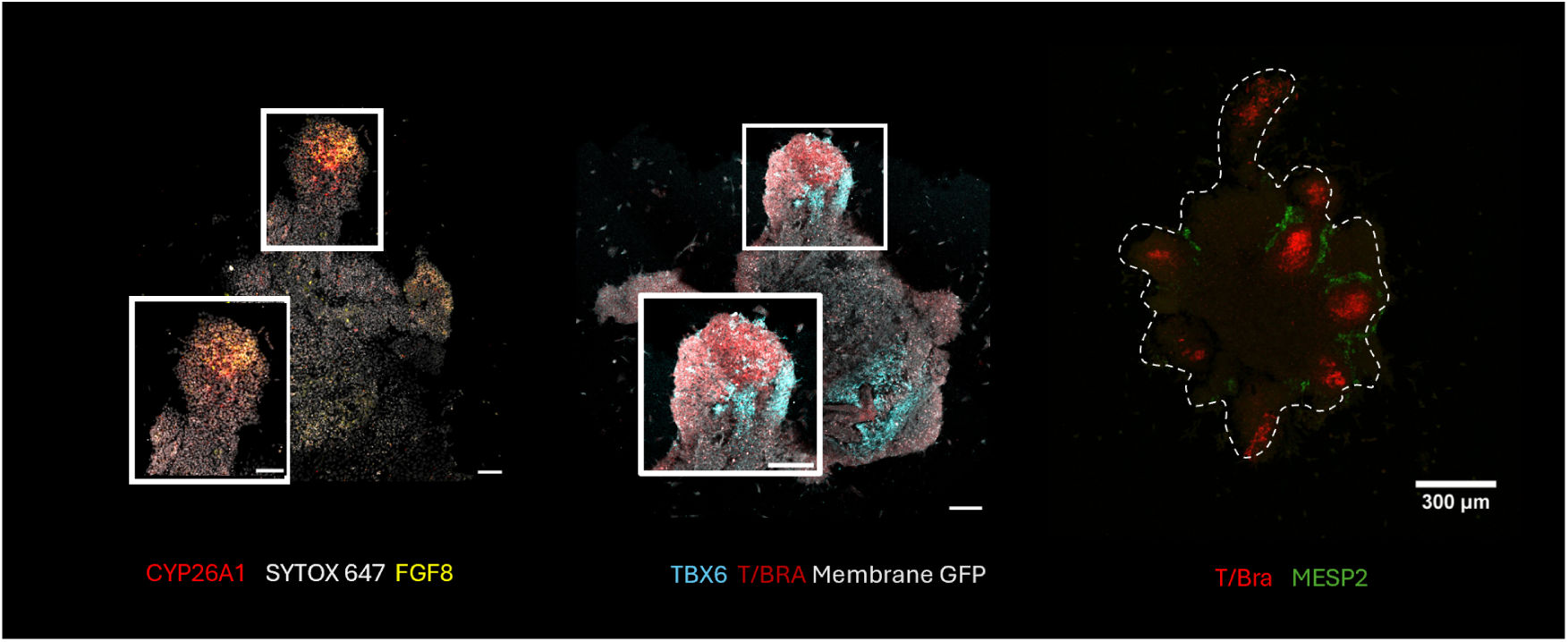
More cell fate markers in gastruloids grown on laminin. A) HCR staining for markers of the posterior tailbud CYP26A1 and FGF8. B) TBX6 (Presomitic mesoderm) and T/Bra (mesoderm). C) MESP2 (Presomitic mesoderm) and T/Bra (mesoderm).

**Supplementary Fig. 5.**
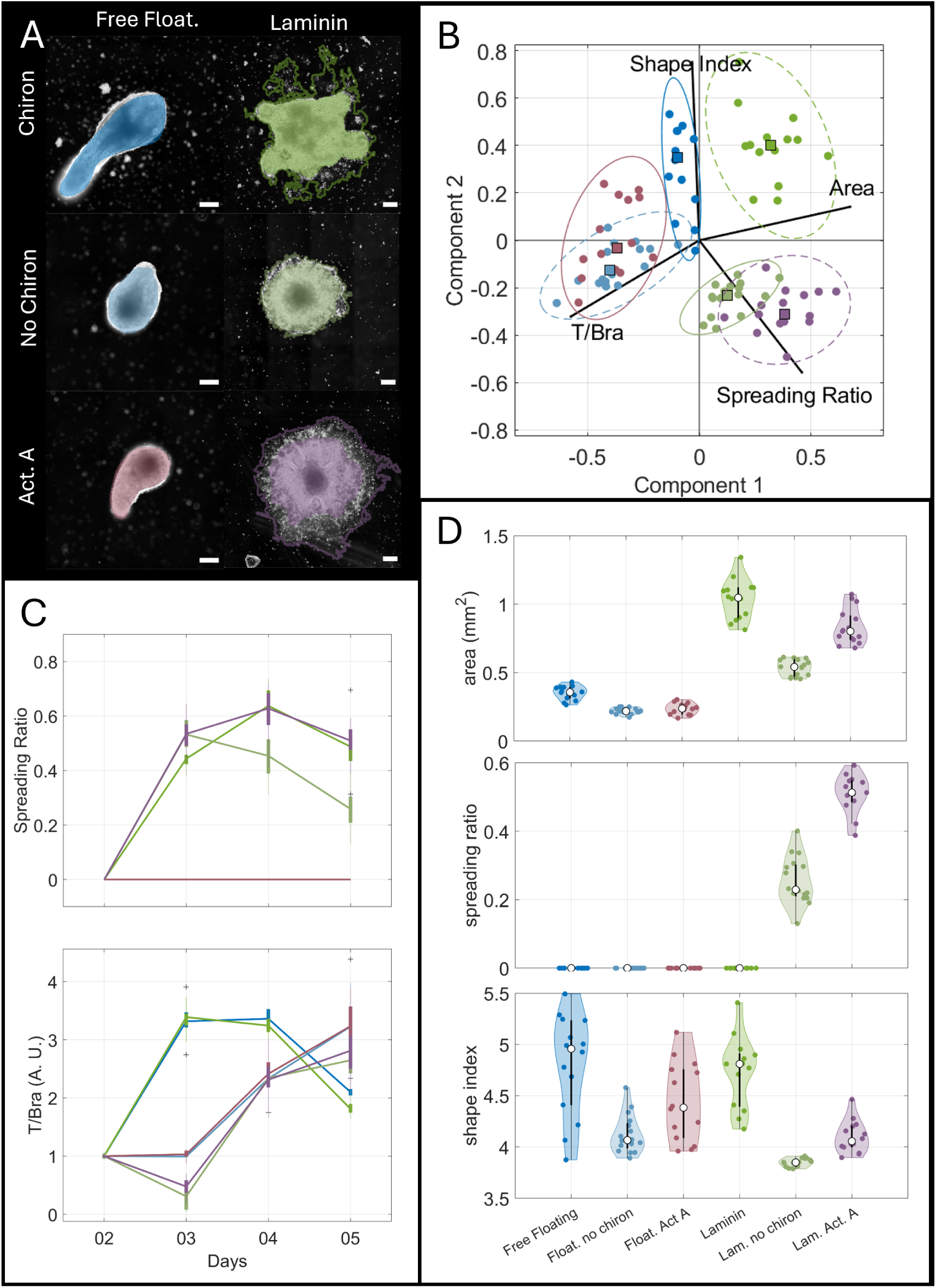
Anterior fates are necessary for cell stream formation. A) Gastruloid morphologies on different substrates with a Chiron pulse or an Activin A pulse on day 2. Only Chiron generates cell streams when seaded on laminin. B) Principal Component Analysis integrating different properties on day 5. C) Analysis of the spreading capability of gastruloids on different substrates (top) and T/Bra expression (bottom) from day 2 to day 5. D) Analysis of area, migratory abilities (spreading ratio) and elongation (shape index) on day 5 gastruloids grown on different coatings in the presence/absence of Chiron. Colours indicate the same conditions across graphs. See the supplementary material for the full statistical analysis.

**Supplementary Fig. 6.**
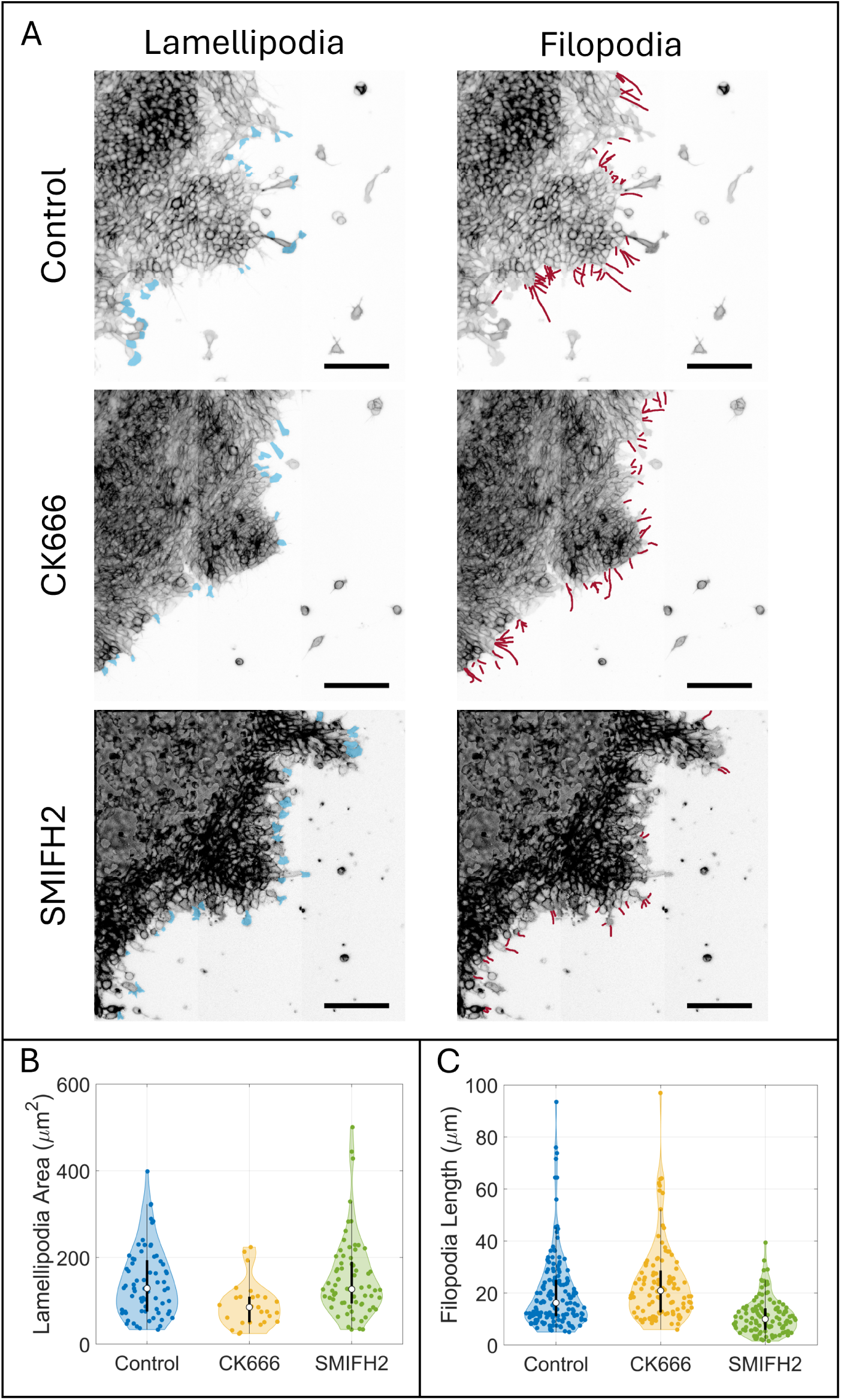
Quantification of Lamellipodia and Filopodia on laminin gastruloids. A) Demonstration of the manual labelling of lamellipodia (left) and filopodia (right) and its morphology with different inhibitors. B) Quantification of lamellipodia area. C) Quantification of filopodia length. See the supplementary material for the full statistical analysis.

**Supplementary Fig. 7.**
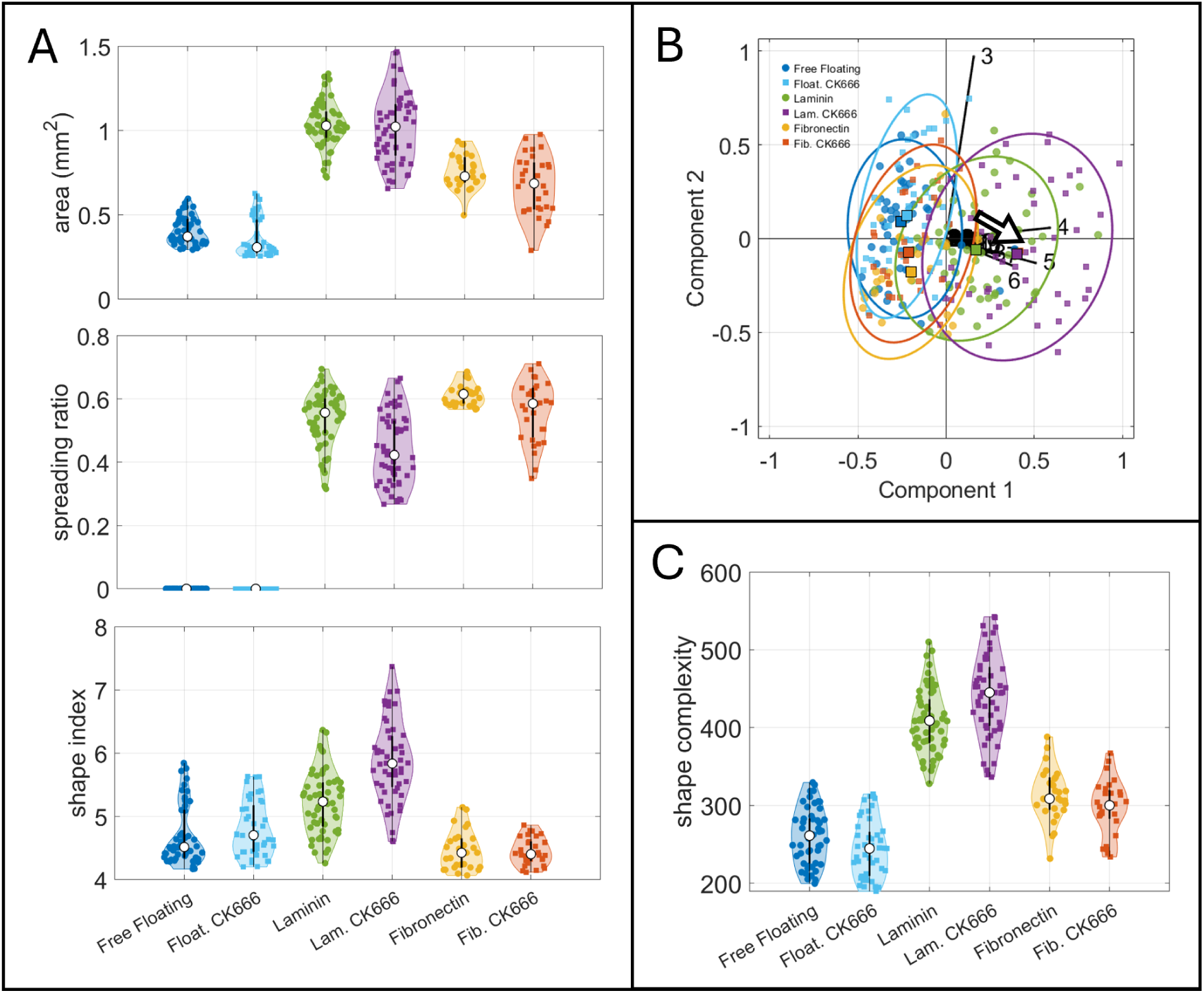
Characterisation of lamellipodia inhibition on different substrate. Analysis of area, migratory abilities (spreading ratio) and elongation (shape index) on day 5 gastruloids grown on different coatings in the presence/absence of CK666 50 *µM*. B) LOCO-EFA modes for the different morphologies. Arrow indicates the need of higher frequency modes to describe the shapes on laminin gastruloids when in presence of the CK666. C) Shape complexity estimated using of the contribution of LOCO-EFA high frequency modes (see methods). See supplementary material for the full statistical analysis.

**Supplementary Fig. 8.**
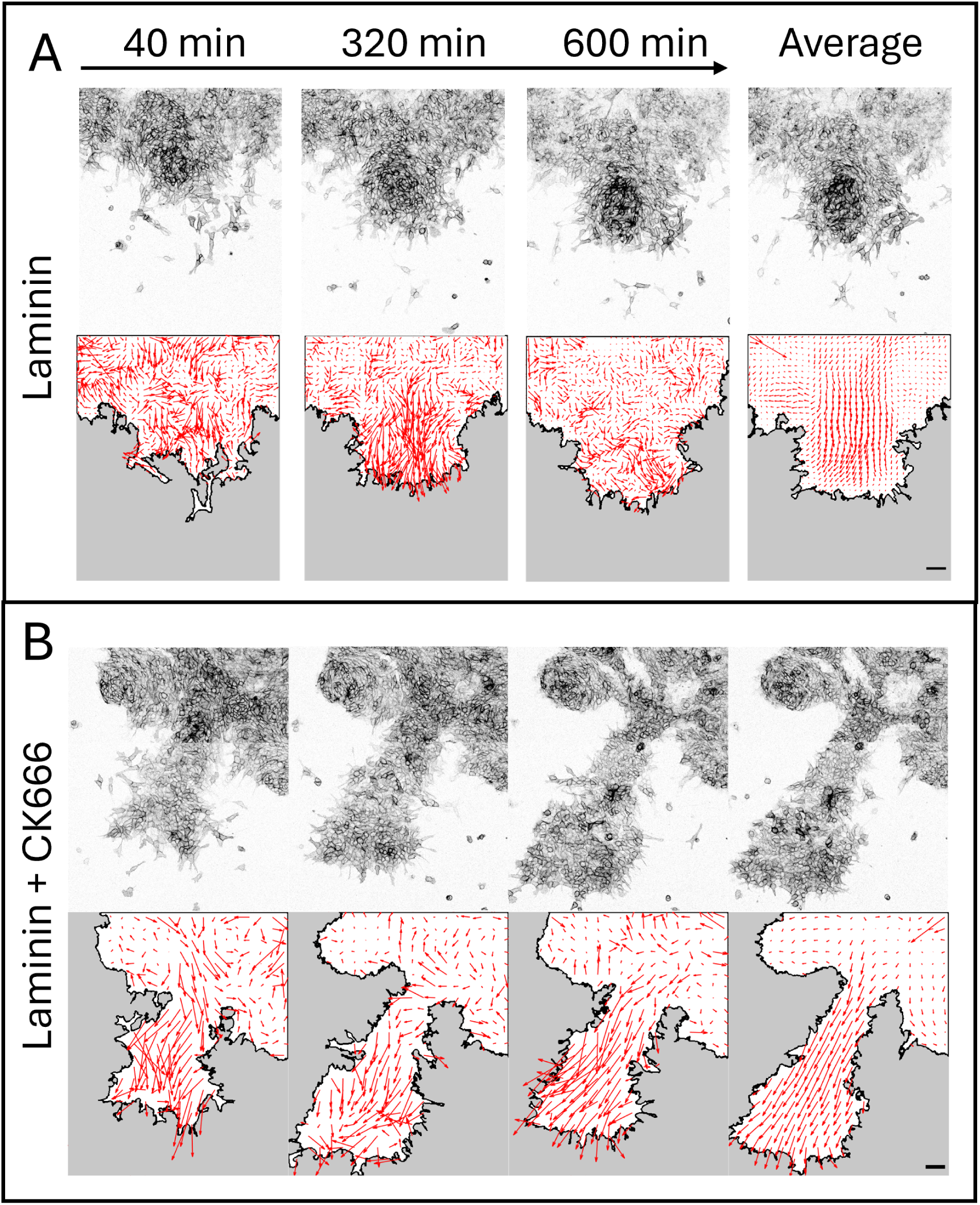
Quantification of tissue flows during cell stream formation. A) Particle Image Velocimetry on laminin and B) in the presence of CK666 50 *µM*. Scale bars 200 *µm*.

**Supplementary Fig. 9.**
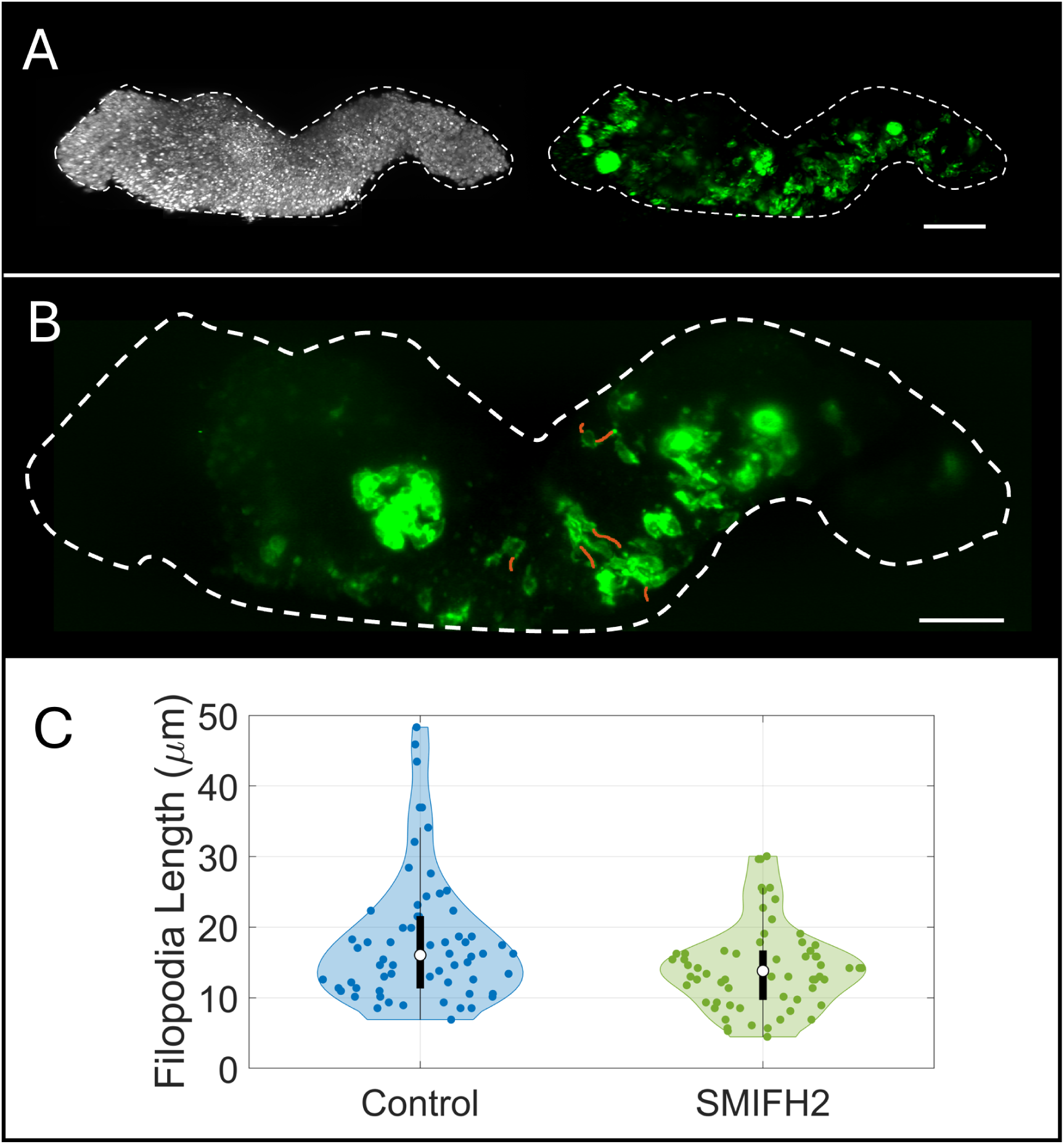
Quantification of Filopodia on free-floating gastruloids. A) Mosaic gastruloid (5% mGFP/ 95% SBN) for quantification of cellular protrusions. B) Demonstration of the manual labelling of filopodia. C) Quantification of filopodia length. See the supplementary material for the full statistical analysis.

**Supplementary Fig. 10.**
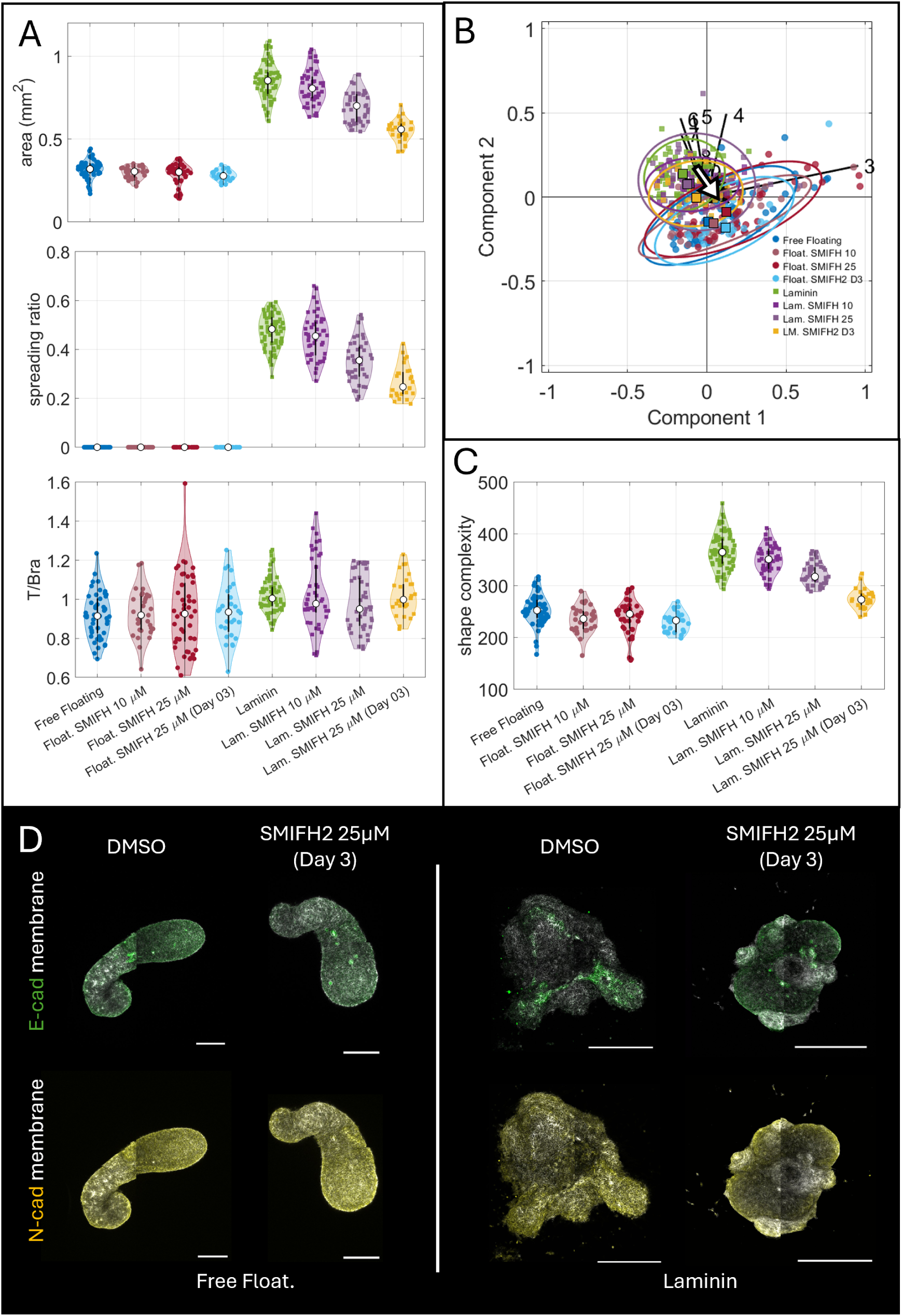
Characterisation of filopodia inhibition on different substrate. Analysis of area, migratory abilities (spreading ratio) and T/Bra expression on day 5 gastruloids, grown free floating or on laminin in the presence/absence of SMIFH2 25 *µM*. B) LOCO-EFA modes for the different morphologies. Arrow indicates the decrease of higher frequency modes to describe the shapes on laminin gastruloids when in presence of the SMIFH2. C) Shape complexity estimated using of the contribution of LOCO-EFA high frequency modes (see methods). D) E-cadh and N-cadh expression on day 5 after in the presence of SMIFH2. See supplementary material for the full statistical analysis.

**Supplementary Fig. 11.**
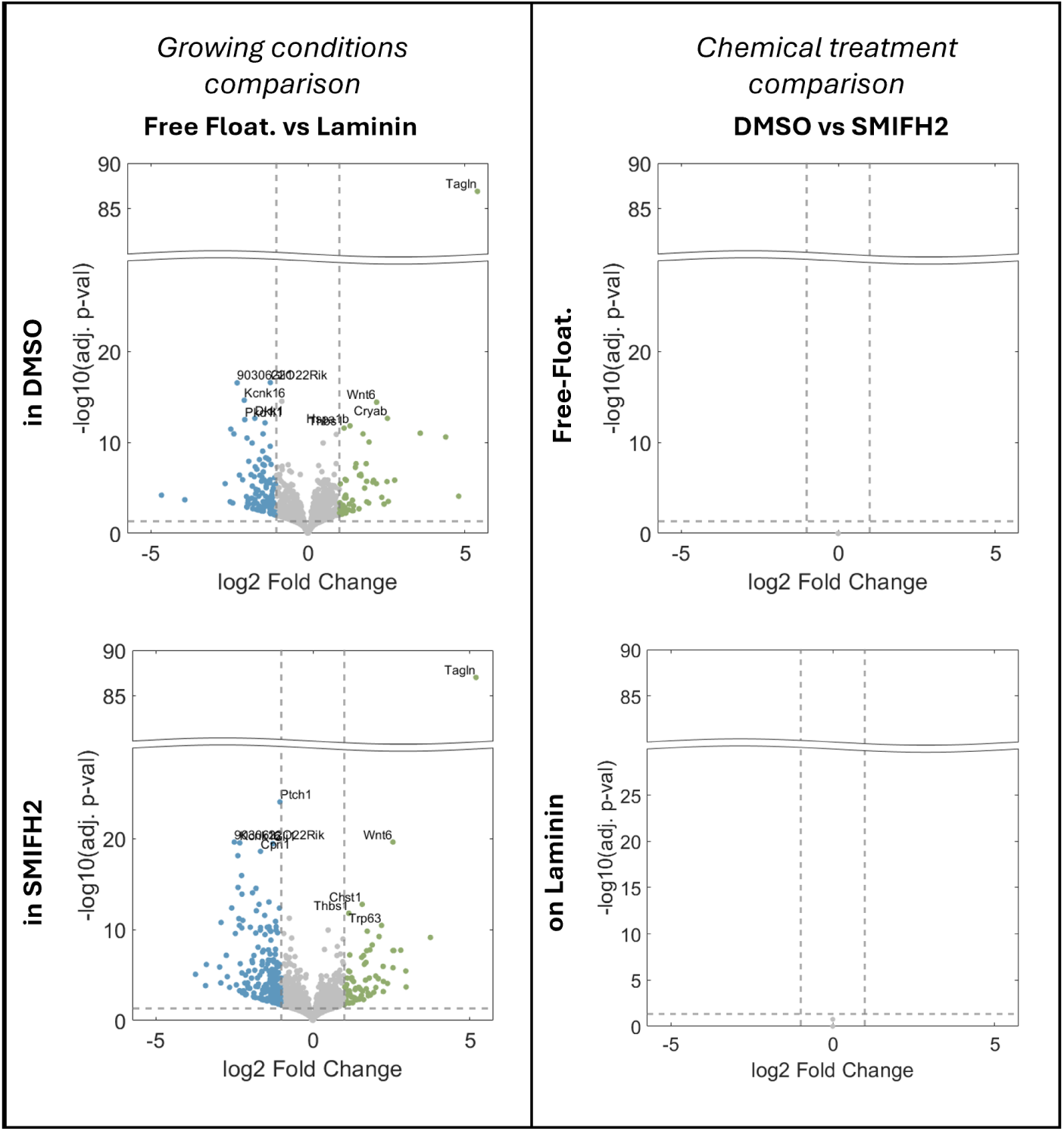
SMIFH2 treatment does not induce transcriptional changes. Volcano plots comparing gene expression across conditions. Left column: comparison between free-floating and laminin gastruloids, either in DMSO (top) or SMIFH2 (bottom). Both conditions show similar patterns of differential gene expression, with genes downregulated on laminin (blue) and upregulated on laminin (green). Right column: comparison between DMSO and SMIFH2 treatment, either in free-floating (top) or laminin (bottom) conditions. No differentially expressed genes are detected in either comparison, confirming that SMIFH2 blocks axis elongation through mechanical effects rather than transcriptional changes. Dashed lines indicate log2FC = ±1 and p-adj = 0.05. Selected genes are labelled.

**Supplementary Fig. 12.**
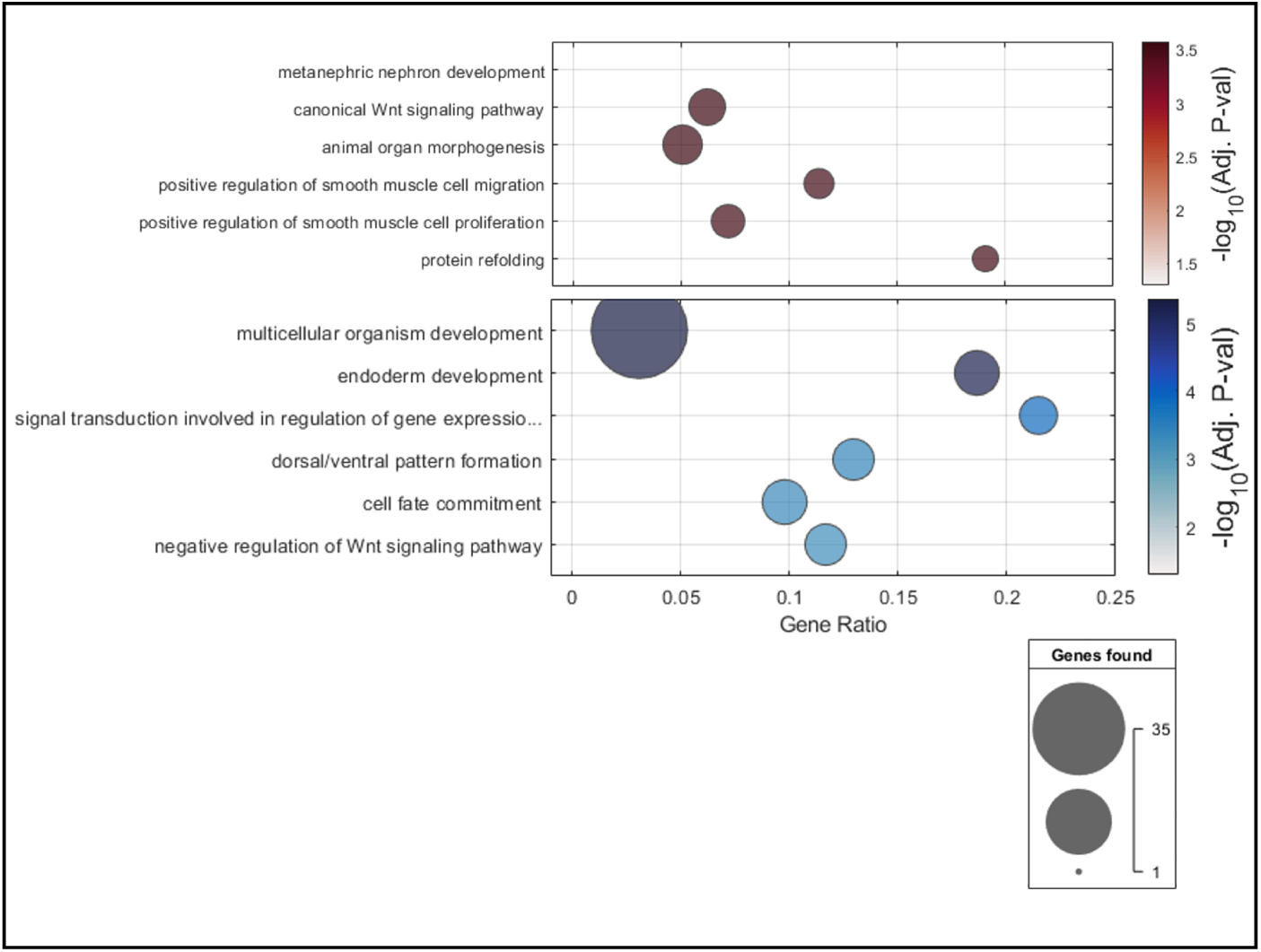
Gene Ontology analysis of differentially expressed genes between laminin and free-floating gastruloids. GO Biological Process enrichment analysis of genes upregulated (top, red) or downregulated (bottom, blue) on laminin compared to free-floating conditions. The x-axis shows the gene ratio (proportion of differentially expressed genes within each GO term), dot size indicates the number of genes found, and colour intensity indicates statistical significance (-log10 adjusted p-value). Only genes with |log2FC| > 1 and p-adj < 0.05 were included in the analysis.

**Supplementary Fig. 13.**
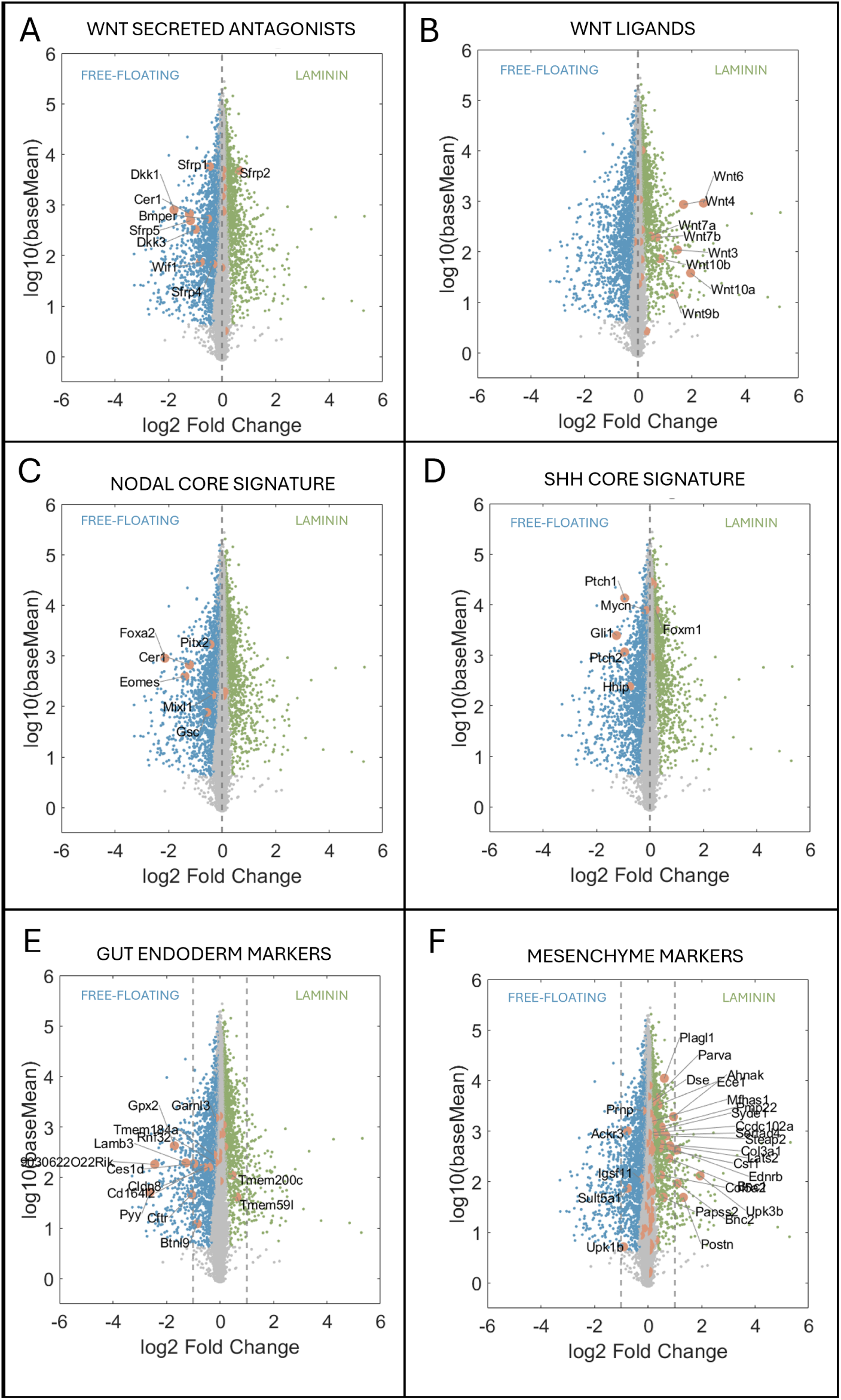
Laminin promotes posterior migratory fates and reduces anterior primitive streak derivatives. Volcano plots highlighting expression levels of specific gene categories comparing laminin versus free-floating gastruloids. A) Wnt secreted antagonists are downregulated on laminin. B) Wnt ligands are upregulated on laminin. C) Nodal pathway core signature genes are downregulated on laminin. D) Shh pathway targets are downregulated on laminin. E) Gut endoderm markers are downregulated on laminin. F) Mesenchyme markers are upregulated on laminin. Together, these data indicate that laminin biases gastruloid differentiation toward posterior, migratory fates while reducing anterior primitive streak derivatives including definitive endoderm. Coloured points indicate genes with p-adj < 0.05; orange points highlight genes of interest within each category. Dashed lines indicate log2FC = ±1.

**Supplementary Fig. 14.**
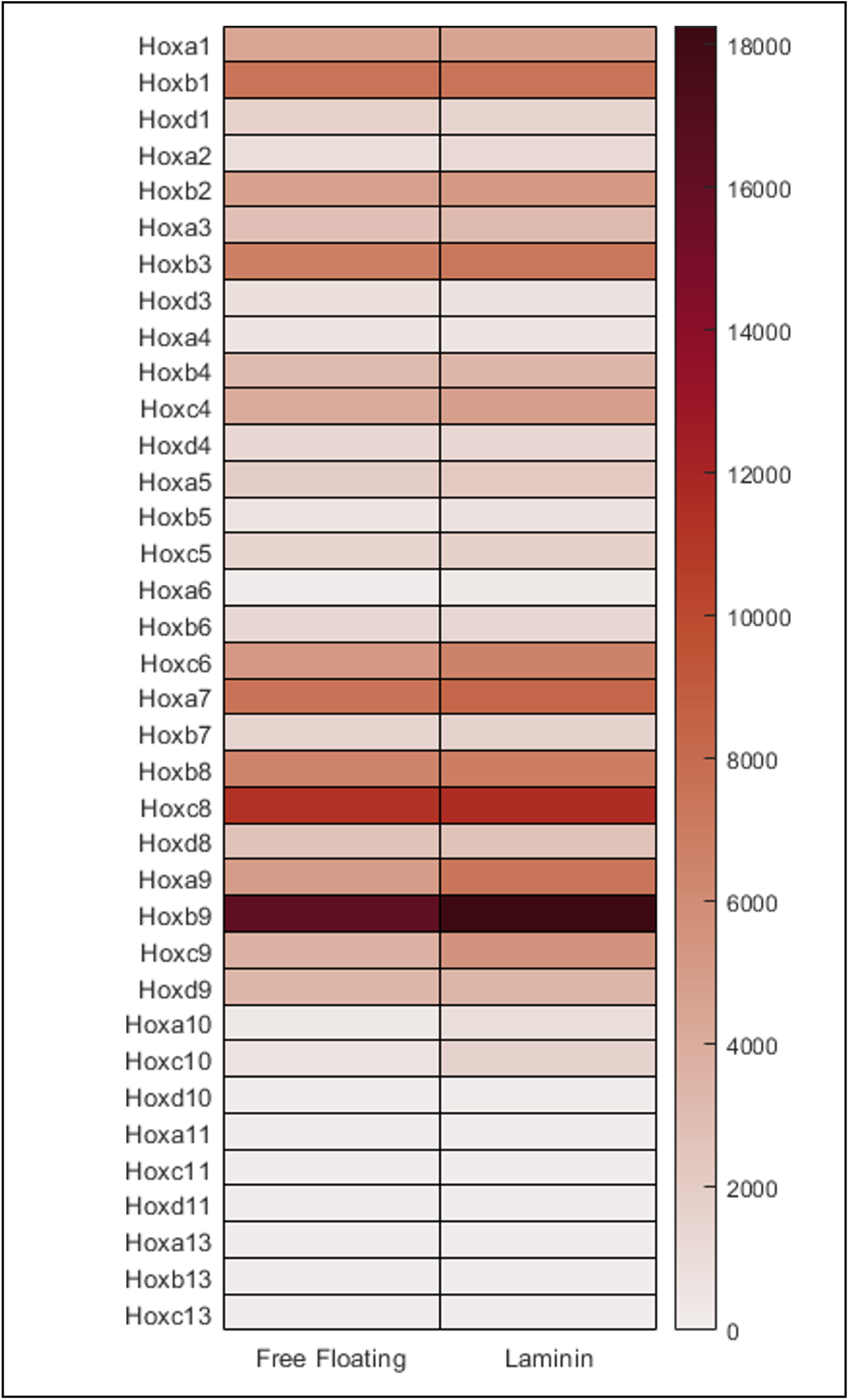
Hox gene expression is unchanged between free-floating and laminin gastruloids. Heatmap showing expression levels of Hox genes in free-floating and laminin gastruloids. Both conditions show similar expression patterns, with high expression of anterior and trunk Hox genes (Hox1-9) and low expression of caudal Hox genes (Hox10-13). This indicates that despite differences in morphogenesis and signalling profiles, both conditions generate tissues with equivalent posterior trunk identity.

**Supplementary Fig. 15.**
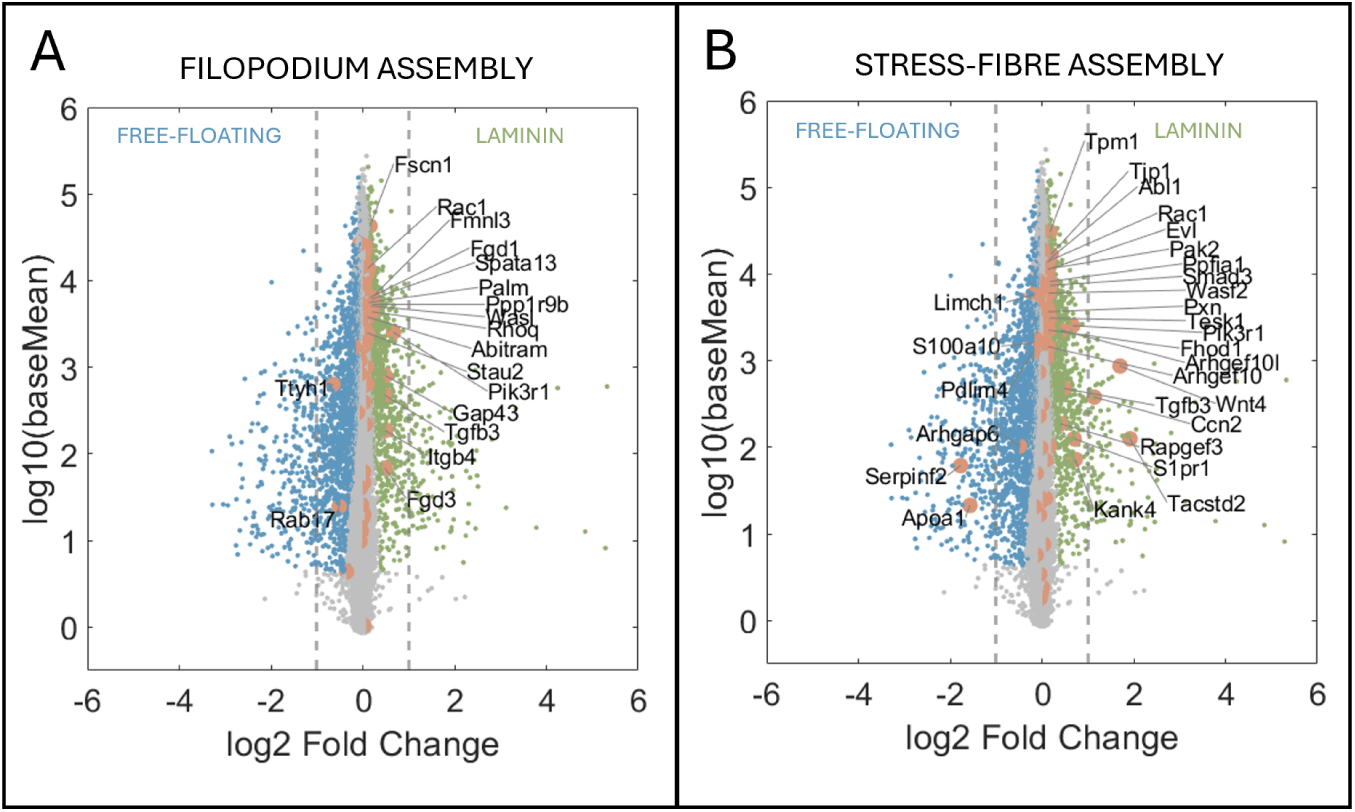
Laminin induces regulators of filopodia and stress-fibres. Volcano plots highlighting expression levels of specific gene categories comparing laminin versus free-floating gastruloids. A) Regulators of filopodium assembly. B) Regulators of stress fibre assembly. Coloured points indicate genes with p-adj < 0.05; orange points highlight genes of interest within each category. Dashed lines indicate log2FC = ±1.

